# Regulation of mitochondrial proteostasis by the proton gradient

**DOI:** 10.1101/2021.12.12.470907

**Authors:** Maria Patron, Daryna Tarasenko, Hendrik Nolte, Mausumi Ghosh, Yohsuke Ohba, Yvonne Lasarzewski, Zeinab Alsadat Ahmadi, Alfredo Cabrera-Orefice, Akinori Eyiama, Tim Kellermann, Elena I. Rugarli, Ulrich Brandt, Michael Meinecke, Thomas Langer

## Abstract

Mitochondria adapt to different energetic demands reshaping their proteome. Mitochondrial proteases are emerging as key regulators of these adaptive processes. Here, we use a multi-proteomic approach to demonstrate regulation of the m-AAA protease AFG3L2 by the mitochondrial proton gradient, coupling mitochondrial protein turnover to the energetic status of mitochondria. We identify TMBIM5 (previously also known as GHITM or MICS1) as a Ca*^2+^*/H*^+^* exchanger in the mitochondrial inner membrane, which binds to and inhibits the m-AAA protease. TMBIM5 ensures cell survival and respiration, allowing Ca*^2+^* efflux from mitochondria and limiting mitochondrial hyperpolarization. Persistent hyperpolarization, however, triggers degradation of TMBIM5 and activation of the m-AAA protease. The m-AAA protease broadly remodels the mitochondrial proteome and mediates the proteolytic breakdown of respiratory complex I to confine ROS production and oxidative damage in hyperpolarized mitochondria. TMBIM5 thus integrates mitochondrial Ca*^2+^* signaling and the energetic status of mitochondria with protein turnover rates to reshape the mitochondrial proteome and adjust the cellular metabolism.

## Introduction

Mitochondria are the major site of cellular ATP production in aerobic organisms and provide numerous metabolites to ensure cell survival, proliferation and differentiation. Mitochondrial integrity depends on multiple intra-organellar proteases, which preserve proteostasis and are emerging as key regulators of mitochondrial plasticity (Deshwal et al., 2020; Song et al., 2021). This is exemplified by *m*-AAA proteases, conserved and ubiquitously expressed ATP dependent metallopeptidases in the inner membrane (IM) of mitochondria (Levytskyy et al., 2017; Patron et al., 2018). m-AAA proteases form homo- or hetero-oligomeric, hexameric complexes, which are built up of homologous AFG3L2- or AFG3L2 and SPG7 (paraplegin) subunits, respectively. Mutations in these subunits are associated with pleiotropic neurodegenerative disorders, ranging from dominant spinocerebellar ataxia (SCA28), recessive hereditary spastic paraplegia (HSP type 7) to dominant optic atrophy (Casari et al., 1998; Charif et al., 2020; Di Bella et al., 2010; Pfeffer et al., 2014; Pierson et al., 2011).

The role of the m-AAA protease for protein quality control was originally recognized in yeast (Arlt et al., 1996; Leonhard et al., 1999) and later shown to be conserved in mammalian cells, where non-assembled and damaged OXPHOS subunits are degraded by the m-AAA protease (Hornig-Do et al., 2012; Zurita Rendon et al., 2014). The loss of m-AAA protease subunits SPG7 or AFG3L2 causes respiratory deficiencies, reduced respiratory complex I activity and increased sensitivity to oxidative stress (Almajan et al., 2012; Atorino et al., 2003; Ehses et al., 2009). Aberrant protein accumulation in the IM reduces the mitochondrial membrane potential (*ΔΨ*m) and causes mitochondrial fragmentation (Ehses et al., 2009; Richter et al., 2015). However, the functional analysis of m-AAA protease deficient cells revealed diverse roles of the m-AAA protease going far beyond its role for protein quality control and the removal of damaged proteins. Processing of MRPL32, a subunit of mitochondrial ribosomes, by the m-AAA protease allows ribosome assembly and mitochondrial translation (Almajan et al., 2012; Nolden et al., 2005). Moreover, the m-AAA protease ensures the assembly of the mitochondrial calcium uniporter (MCU) complex in the IM by degrading excess EMRE subunits (Konig et al., 2016; Tsai et al., 2017). This prevents the formation of MCU complexes that lack the regulatory subunits MICU1 and MICU2, counteracting uncontrolled Ca^2+^ influx into mitochondria, mitochondrial Ca^2+^ overload and cell death. The m-AAA protease thus maintains mitochondrial Ca^2+^ homeostasis and ensures cell survival (Patron et al., 2018), which may also be of relevance for neurodegeneration (Maltecca et al., 2015).

Although studies in various mouse models and cultured cells revealed critical functions of m-AAA proteases within mitochondria, it remained enigmatic how the m-AAA protease activity is regulated. Here, we identify TMBIM5 (Transmembrane BAX Inhibitor-1 Motif 5; previously also known as GHITM or MICS1 (Li et al., 2001; Oka et al., 2008)) as a novel interactor and inhibitor of the m-AAA protease. We demonstrate that TMBIM5 acts as a Ca^2+^/H^+^ exchanger in the IM, which limits mitochondrial hyperpolarization and ROS production and couples the activity of the m-AAA protease to the energetic demands of the cell. Persistent hyperpolarization causes degradation of TMBIM5 and leads to complex I degradation and broad reshaping of the mitochondrial proteome by the m-AAA protease.

## Results

### Identification of TMBIM5 as a novel interactor of the m-AAA protease

In an attempt to identify interacting proteins and substrates of human AFG3L2, we immunoprecipitated AFG3L2 from mitochondria of Flp-In HEK293 T-REx cells expressing proteolytically inactive AFG3L2^E408^-FLAG. The analysis of co-purifying proteins by mass spectrometry identified paraplegin (SPG7), a proteolytic subunit of the *m*-AAA protease complex, and known *m*-AAA protease binding proteins, including the prohibitin membrane scaffold complex (PHB1, PHB2) and MAIP1 (Figure 1A, Table S1). Besides these expected interactors, TMBIM5 (also known as GHITM or MICS1) was highly enriched in AFG3L2 precipitates (Figure 1A, Table S1), which was further validated by immunoblot analysis (Figure S1A). These experiments identified TMBIM5 as a novel interactor of the *m*-AAA protease.

**Figure 1.**
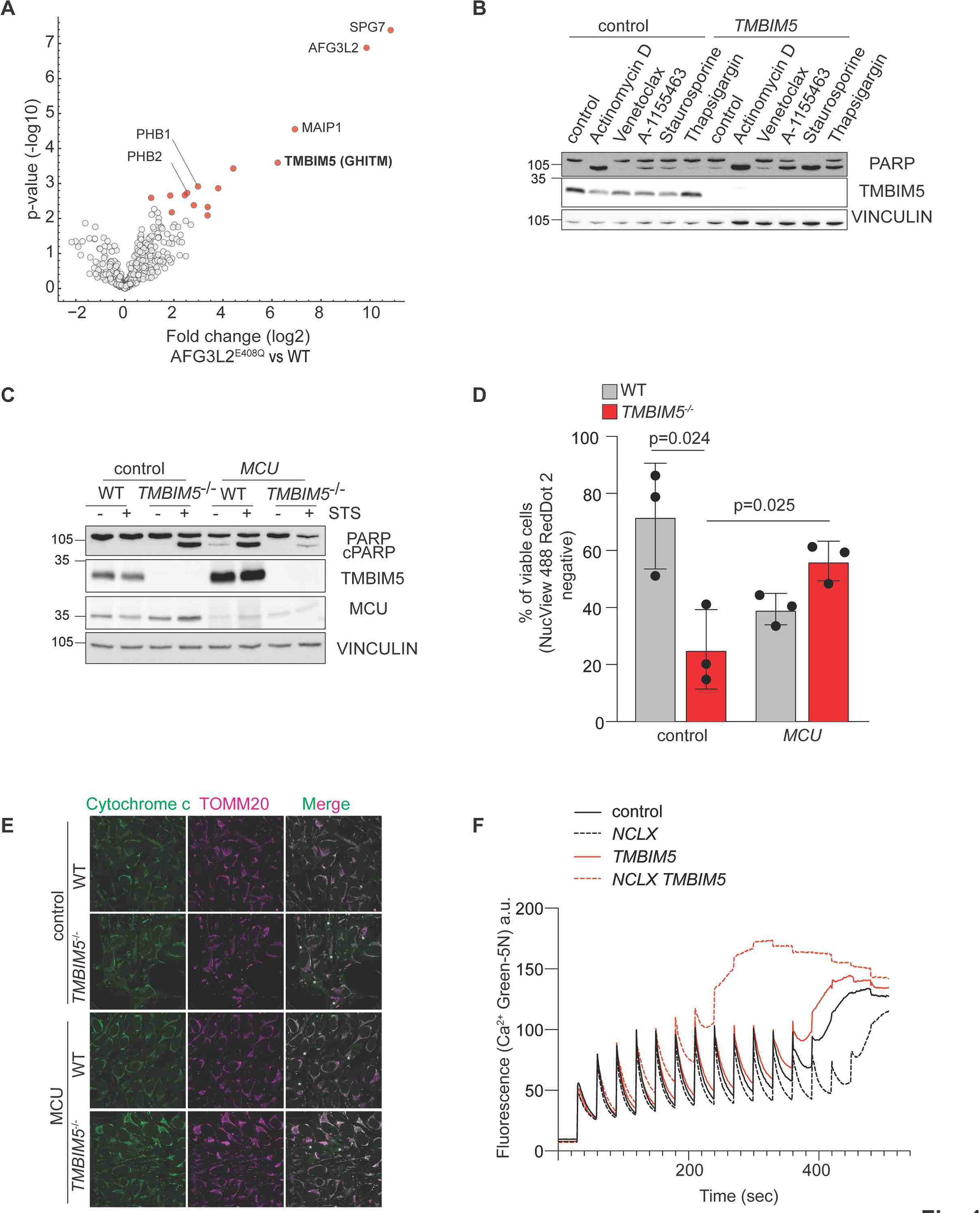
Mitochondrial Ca*^2+^* overload triggers apoptotic death of *TMBIM5^-/-^* cells. (A) Volcano plot representation of AFG3L2^E408Q^-FLAG interaction partners. Mitochondria isolated from Flp-In HEK293T-REx cells expressing AFG3L2^E408Q^-FLAG were subjected to immunoprecipitation. Co-purifying proteins were identified by quantitative mass spectrometry (MS) (n=5 independent experiments) followed by an unpaired two-sided t-test and permutation-based FDR estimation to correct for multiple testing (FDR < 0.05. Significantly enriched MitoCarta 3.0 proteins (Rath et al., 2021) are colored in red, gene names indicate previously verified AFG3L2 interactors and TMBIM5 (GHITM, MICS1). See also Table S1. (B) Representative immunoblot of HeLa wildtype (WT) cells transfected with scrambled siRNA (control) or siRNA targeting *TMBIM5* and treated with the indicated drugs for 16 h. Control (0.1% DMSO), actinomycin D (1 µM), venetoclax (1 µM), A-1155463 (1 µM), staurosporine (1 µM), thapsigargin (2 µM). Quantification is shown in Figure S1B (n=3 independent experiments). (C) Representative immunoblot of HeLa WT and *TMBIM5*^-/-^ cells transfected with scrambled siRNA (control) or siRNA targeting *MCU* for 72 h. Where indicated samples were treated with staurosporine (STS; 1 µM) for 16 h. Quantification is shown in Figure S1I (n=3 independent experiments). (D) Viability of WT and *TMBIM5*^-/-^ cells after incubation with staurosporine (STS; 0.1 µM) for 16 h. Cell viability was assessed cytofluorimetrically monitoring NucView488 and RedDot2 staining and expressed as percentage of all cells. (n=3 independent experiments). Two-tailed *t*-test. A p-value of <0.05 was considered statistically significant. (E) WT and *TMBIM5*^-/-^ HeLa cells were transfected with the indicated scrambled siRNA (control) and siRNA targeting *MCU* for 48 h and incubated with staurosporine (0.1 µM) for 16 h in the presence of Z-VAD-FMK (50 µM) and epoxomicin (1 µM) to prevent apoptosis. Cells were subjected to immunofluorescence analysis with antibodies directed against cytochrome c (green) and TOMM20 (magenta). *Cells with cytosolic cytochrome *c*. Quantification is shown in Figure S1L (n=3 independent experiments; number of cells > 150 each experiment). (F) Calcium retention capacity (CRC) was assessed in isolated mitochondria of HEK293T WT cells transfected with the indicated siRNA. The experiment was performed in sucrose-based medium containing respiratory substrates and the membrane-impermeable Ca^2+^ sensor Ca^2+^ Green-5N. Ca^2+^ Green-5N fluorescence was monitored following repeated addition of Ca^2+^ pulses (10 µM). Data show representative traces (2 independent experiments run in triplicate each condition). See also Figure S1 and Table S1.

### Mitochondrial Ca^2+^ overload triggers apoptotic death of TMBIM5^-/-^ cells

TMBIM5 belongs to an evolutionary conserved family of six membrane proteins (TMBIM1- 6), which contain the transmembrane BAX inhibitor motif, originally described for the founding member of this family BI-1 (or TMBIM6). TMBIM5 was originally identified as a deregulated protein upon growth factor withdrawal in mice (Li *et al*., 2001) and recently found in CRISPR/Cas9 screens to be essential in human cells cultivated on physiological medium (Rossiter et al., 2021). In contrast to other TMBIM proteins, TMBIM5 is localized to the IM, where it protects against apoptosis (Oka et al., 2008; Seitaj et al., 2020). In agreement with these findings, we observed an increased susceptibility of HeLa cells depleted of TMBIM5 to different apoptotic stimuli, when monitoring PARP cleavage (Figure 1B). Similarly, *TMBIM5*^- /-^ HeLa cells, which were generated by CRISPR/Cas9-mediated genome editing, were more sensitive to apoptotic cell death induced by staurosporine (Figure S1B), as has been observed in TMBIM5-deficient HAP1 cells (Seitaj et al., 2020). It was suggested that TMBIM5 preserves cristae morphogenesis and interacts with cytochrome *c*, limiting its release into the cytoplasm in apoptotic cells (Oka et al. 2008). However, *TMBIM5*^-/-^ HeLa cells harbored a normal, reticulated mitochondrial network with unaltered cristae (Figure S1C-F), suggesting that other mechanisms lead to the increased apoptotic vulnerability of *TMBIM5*^-/-^ cells.

Members of the TMBIM family function as pH-dependent Ca^2+^ leak channels and regulate both Ca^2+^ homeostasis and cell death (Kim et al., 2008; Liu, 2017; Rojas-Rivera and Hetz, 2015). To investigate the possibility that altered mitochondrial Ca^2+^ homeostasis causes the increased vulnerability of *TMBIM5*^-/-^ cells, we monitored PARP cleavage as readout for cell death in these cells after silencing the mitochondrial calcium uniporter, MCU. Depletion of MCU from *TMBIM5*^-/-^ cells reduced PARP cleavage upon treatment with staurosporine (Figure 1C). Moreover, MCU depletion significantly reduced the number of apoptotic or necrotic *TMBIM5*^-/-^ cells harboring cleaved caspase 3/7 (Figure 1D, Figure S1G) and the release of cytochrome *c* from TMBIM5-deficient mitochondria to the cytosol (Figure 1E, Figure S1H). Therefore, limiting MCU-dependent mitochondrial Ca^2+^ uptake protects *TMBIM5*^-/-^ cells against apoptosis, suggesting that cell death is induced by mitochondrial Ca^2+^ overload.

To corroborate these findings, we determined the Ca^2+^ retention capacity (CRC) of mitochondria depleted of TMBIM5, which were isolated from HEK293T cells. To this end, we monitored Ca^2+^ influx after repeated Ca^2+^ pulses using the Ca^2+^-dependent fluorescent indicator Calcium-Green 5N (Figure 1F). The loss of TMBIM5 alone did not significantly affect the CRC, but resulted in lowered CRC and earlier opening of the membrane permeability transition pore (MPTP) upon additional depletion of the mitochondrial Na^+^/Ca^2+^ exchanger NCLX (Figure 1F). Thus, HEK293T cells depleted of TMBIM5 are more susceptible to mitochondrial Ca^2+^ overload. Ca^2+^ export by NCLX can at least partially compensate for the loss of TMBIM5, indicating that loss of TMBIM5 reduces Ca^2+^ leakage from mitochondria rather than increasing Ca^2+^ influx.

### TMBIM5 acts as a Ca^2+^/H^+^ exchanger in the inner membrane

To further investigate the role of TMBIM5 as a potential mitochondrial ion channel, we measured mitochondrial Ca^2+^ uptake after stimulating HeLa cells with thapsigargin, an inhibitor of the sarco-endoplasmic reticulum calcium ATPase (SERCA). Thapsigargin treatment induces Ca^2+^ leakage from the ER and promotes mitochondrial Ca^2+^ uptake at ER-mitochondria contact sites. We expressed an RFP-based Ca^2+^ indicator that was specifically targeted to the mitochondrial matrix (mtRed-GECO; (Wu et al., 2013)) in HeLa cells depleted of TMBIM5 and monitored fluorescence intensity after thapsigargin treatment (Figure 2A). TMBIM5 depleted cells accumulated Ca^2+^ within mitochondria when compared to control cells (Figure 2A), which is consistent with our hypothesis of reduced mitochondrial Ca^2+^ efflux in the absence of TMBIM5.

**Figure 2.**
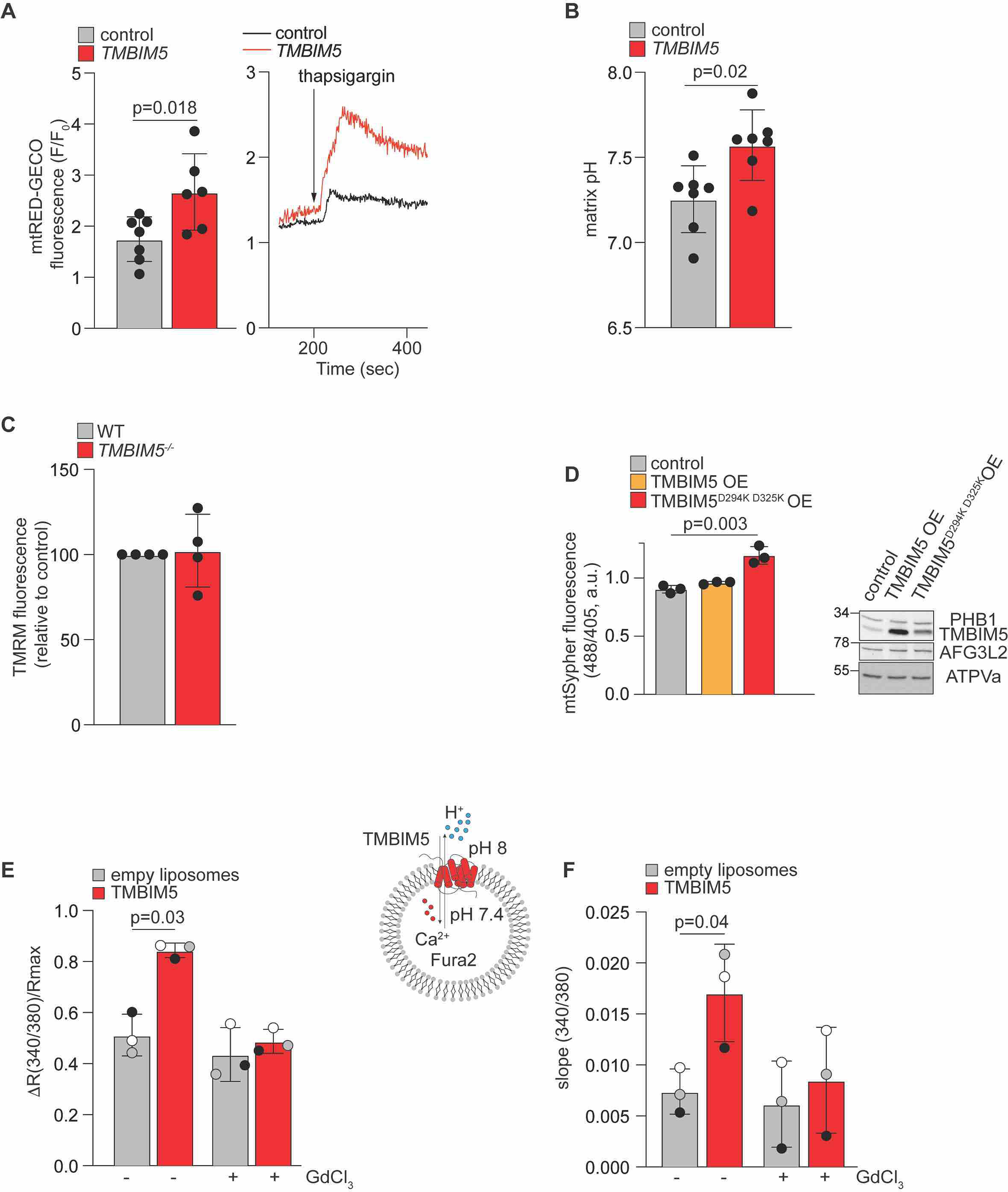
TMBIM5 acts as a Ca*^2+^*/H*^+^* exchanger. **(A)** Fluorescence (F) F/F0 ratio quantifications, where F0 is the average of the first ten seconds and F is the fluorescence peak measured after thapsigargin (2 µM) administration (left panel). The right shows mtRED-GECO ratio traces of HeLa cells transfected with scrambled siRNA (control) and siRNA targeting *TMBIM5*. Each measurement was performed in at least 10 cells from 6 different preparations. Two-tailed *t*-test. A p-value of <0.05 was considered statistically significant. **(B)** Matrix pH was measured with Sypher3smito in HeLa WT transfected with scrambled siRNA (control) or siRNA targeting *TMBIM5* for 72 h. The probe was calibrated in KRB buffer by the stepwise addition/removal of permeant weak acid. Each measurement was performed in at least 80 cells from 7 different preparations. Two-tailed *t*-test. A p-value of <0.05 was considered statistically significant. **(C)** Mitochondrial membrane potential was monitored with TMRM in WT and *TMBIM5*^-/-^ HeLa cells. Fluorescent intensity was calculated as ratio to the value upon CCCP addition (15 µM). Values are expressed as mean relative to control. Each measurement was performed in triplicate from 4 different preparations. **(D)** Matrix pH was measured with Sypher3smito in WT HeLa cells transiently expressing TMBIM5 or TMBIM5^D294K D325K^ for 24 h. The ratio of absorbance at 488 nm and 405 nm is shown in arbitrary units (a.u.). Each measurement was performed in at least 20 cells from 3 different preparations. Representative immunoblot of WT HeLa cells transfected with TMBIM5 or TMBIM5^D294K D325K^ for 24 h (right panel). Two-tailed *t*-test. A p-value of <0.05 was considered statistically significant. **(E)** Liposome assay. Recombinant TMBIM5 was reconstituted in liposomes containing 100 µM Fura-2. CaCl2 was added later at the concentration of 100 µM. Rmax was obtained at the end of the experiment after the addition of ionomycin (10 µM). Data are expressed as mean of R/Rmax, where R is the peak after CaCl2 addition. Where indicated GdCl3 (100 µM) was added 10 min before the experiment. (n=3 independent experiments). Two- tailed *t*-test. A P value of <0.05 was considered statistically significant. **(F)** Liposome assay (as in E). Slope was calculated in the 100 s after 100 µM CaCl2 addition as expressed as mean ±SD. Two-tailed *t*-test. A p-value of <0.05 was considered statistically significant. See also Figure S2.

The founding member of the TMBIM family, TMBIM6 (or BI-1) was suggested to be a pH-sensitive Ca^2+^ leak channel in the ER membrane, which can be activated by cytosolic protonation of a di-aspartyl moiety (Bultynck et al., 2014; Chang et al., 2014; Guo et al., 2019; Li et al., 2020). A Ca^2+^/H^+^ antiporter activity of TMBIM6 has been discussed controversially, mainly because a Ca^2+^/H^+^ exchanger can function only in the presence of a stable H^+^ gradient that is not present between the ER lumen and the cytosol. Since the di-aspartyl pH sensor motif of TMBIM6 is conserved in TMBIM5 and since a H^+^ gradient is constantly present across the IM, we examined whether TMBIM5 can act as a Ca^2+^/H^+^ exchanger. We expressed a mitochondrially targeted variant of the synthetic pH sensor SypHer (SypHer3s-dmito, (Ermakova et al., 2018)) in cells depleted of TMBIM5 and monitored the pH in the mitochondrial matrix (Figure 2B). Strikingly, loss of TMBIM5 increased the matrix pH from pH 7.25 to pH 7.6 (Figure 2B), indicating that the TMBIM5-mediated Ca^2+^ release from mitochondria is indeed accompanied by H^+^ transport into the mitochondrial matrix. TMRM staining showed normal *ΔΨ*m in *TMBIM5*^-/-^ cells when compared to wildtype cells, excluding that the decreased matrix H^+^ content reflects differences in *ΔΨ*m (Figure 2C, S2A) (Chang et al., 2014; Guo et al., 2019). While transient expression of TMBIM5 in HeLa cells did not alter matrix pH, we observed an increased matrix pH when compared to wildtype cells upon expression of TMBIM5^D294K D325K^, harboring mutations in the di-aspartyl motif (Figure 2D). Thus, expression of TMBIM5^D294K D325K^ mimics the phenotype of *TMBIM5*^-/-^ cells, highlighting the importance of the di-aspartyl motif for TMBIM5 function.

To unambiguously demonstrate channel activity of TMBIM5, we purified TMBIM5 expressed in *Escherichia coli* and reconstituted the protein in liposomes harboring the membrane-impermeable Ca^2+^ sensor Fura-2 (Figure 2E; Figure S2C-E). After addition of Ca^2+^ to liposomes, we observed increased Ca^2+^ influx into liposomes containing TMBIM5 when compared to control liposomes in the presence of a pH gradient (Figure 2E, F). Gadolinium chloride (GdCl3), a potent inhibitor of voltage-gated Ca^2+^ channels, efficiently inhibited the TMBIM5-dependent Ca^2+^ influx (Figure 2E, F). Thus, recombinant TMBIM5 is able to facilitate specifically the influx of Ca^2+^ into liposomes.

We then asked whether we can directly show channel activity of TMBIM5. We fused TMBIM5 containing proteo-liposomes with planar-lipid membranes to perform high-resolution electrophysiology measurements (Figure 3). Fusion events were observed and led to stable insertion of channel proteins into planar-lipid bilayers (Figure 3A). From a conductance state analysis based on single gating events a main conductance state of ∼ 50 pS (Figure S3A, B) was found at 250 mM CaCl2. Considering a cylindrical pore with a restriction zone of ∼1 nm in length as the structural basis and assuming a five-fold higher solution resistance within the pore than in the bulk medium as described before (Smart et al., 1997), this corresponds to a channel diameter of ∼0.5 nm. Occasionally, we observed larger gating events, suggesting broader dynamics in gating and pore size. The calculated pore size is in good agreement with the channel diameter at the restriction zone observed in crystal structures of a bacterial homologue of TMBIM5 (Chang et al., 2014). Interestingly, even when titrating down the protein to lipid ratio in proteoliposomes to be fused with planar-lipid membrane, we were only able to measure multiple copies of the channel (Figure 3B), suggesting that TMBIM5 forms oligomers after reconstitution. As the channel activity was stable over a long period of time, we could monitor pore activity upon addition of GdCl3 which inhibits Ca^2+^ uptake into liposomes. GdCl3 let to a significantly decreased conductivity (Figure 3C) and a tendency for gating towards the closed state at elevated applied voltages (Figure 3B, C). At holding potentials of +80 mV the channel displayed a 70% decreased open probability (Figure 3D), showing that calcium conductivity can be blocked by GdCl3.

**Figure 3.**
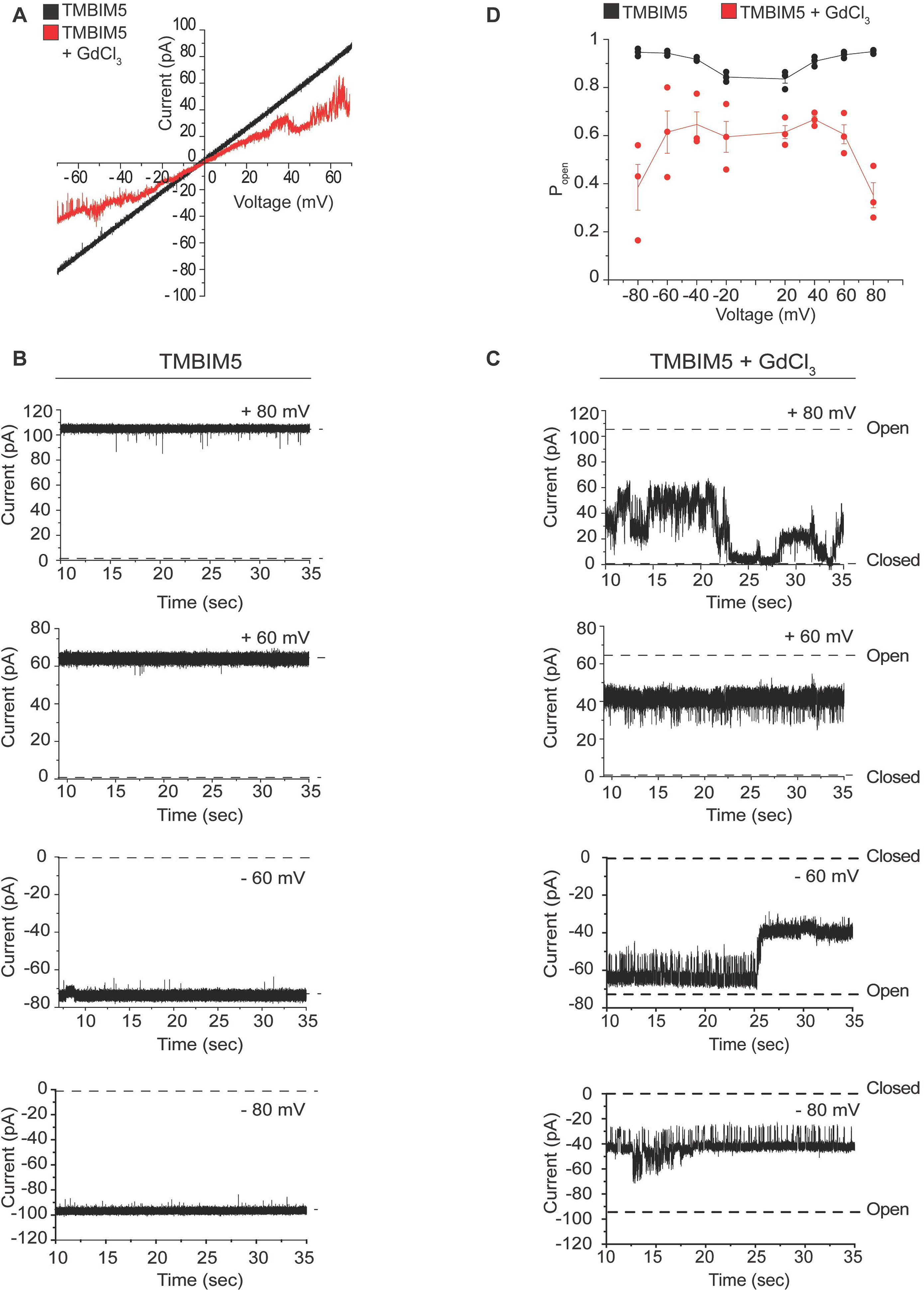
Reconstitution of TMBIM5 activity in lipid bilayers. **(A)** Current-voltage relationship of multiple TMBIM5 channels in the absence (black) and presence (red) of GdCl3. **(B)** Representative current traces of multiple TMBIM5 channels inserted into planar lipid bilayers at indicated holding potentials. **(C)** Representative current traces of the same TMBIM5 channels inserted into planar lipid bilayers in the presence of GdCl3 at indicated holding potentials. Fast gating events and channel closure at elevated membrane potentials can be observed in the presence of GdCl3. **(D)** Voltage-dependent open probability (Popen) of TMBIM5 in the absence and presence of GdCl3. (n=3 independent channel incorporations). See also Figure S3.

Together, we conclude that TMBIM5 can act as a Ca^2+^/H^+^ exchanger in the IM, coupling mitochondrial export of Ca^2+^ with H^+^ import into mitochondria. Loss of TMBIM5 leads to matrix alkalinization, mitochondrial hyperpolarization and the accumulation of Ca^2+^ within mitochondria, rendering cells sensitive to mitochondrial Ca^2+^ overload and cell death.

### TMBIM5 controls oxidative phosphorylation

We determined in further experiments how TMBIM5 affects mitochondrial bioenergetics. *TMBIM5*^-/-^ cells showed decreased basal and maximal oxygen consumption rates and a reduced spare respiratory capacity (Figure 4A-C; Figure S4A-C), as has been observed in TMBIM5- deficient HAP1 cells (Seitaj et al., 2020). The production of ATP was also diminished in these cells (Figure 4D; Figure S4D). Reduced respiration was accompanied by an increased ROS production in *TMBIM5*^-/-^ cells (Figure 4E). When analyzing specific enzymatic activities of individual respiratory chain complexes in isolated mitochondria, we observed a reduced complex I activity, while the activity of the other respiratory complexes was not significantly affected (Figure 4F). These experiments suggest that a reduced specific activity of complex I limits mitochondrial respiration in *TMBIM5*^-/-^ mitochondria.

**Figure 4.**
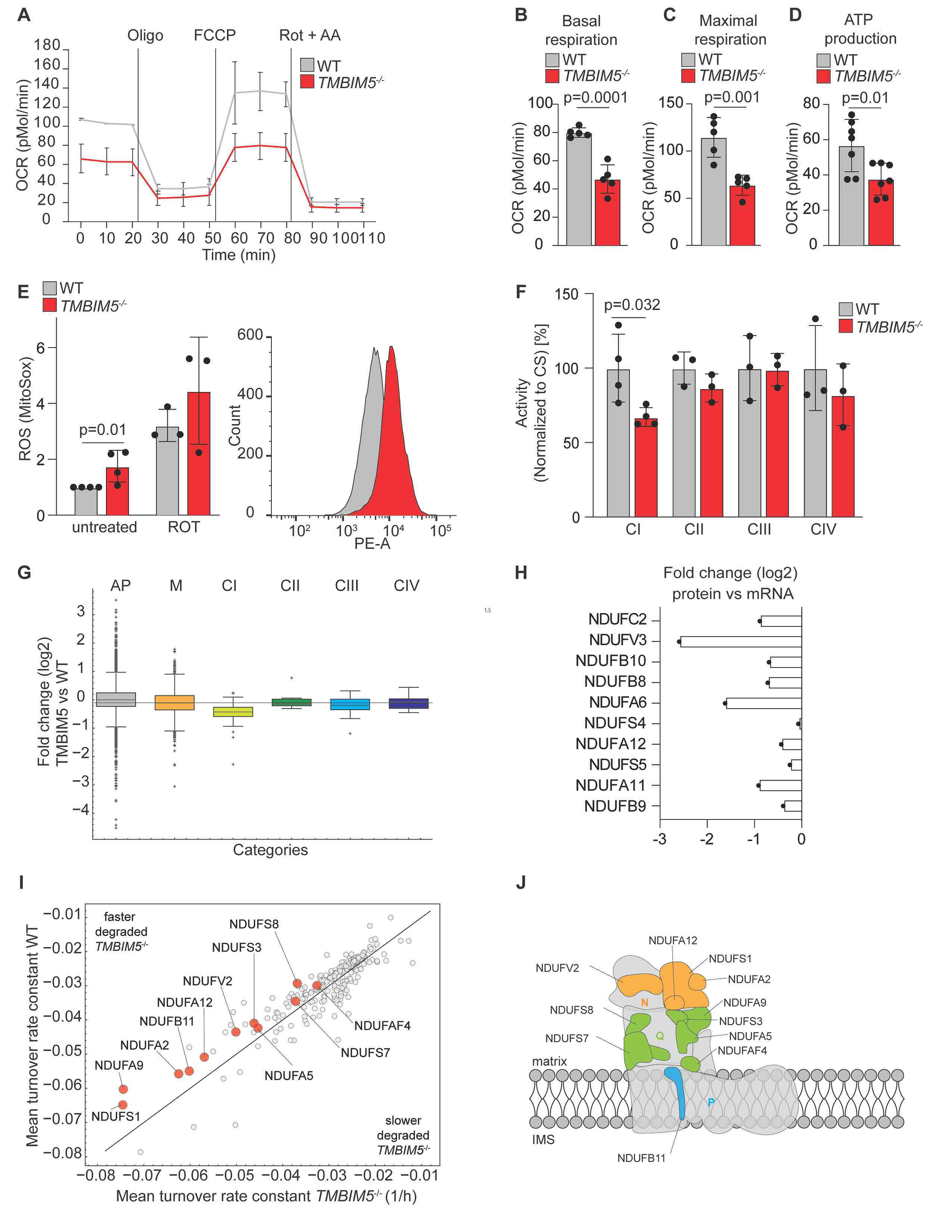
TMBIM5 controls oxidative phosphorylation. **(A)** Oxygen consumption rate (OCR) of WT and *TMBIM5*^-/-^ HeLa cells in glucose-containing media. Traces are mean ± SD of 7 independent experiments each one run at least in triplicate. Labeled lines denotes injections of oligomycin (Oligo, 2 µM), FCCP (0.5 µM), rotenone and antimycin A (Rot + AA; both 0.5 µM). **(B)** Basal respiration calculated from OCR experiment of WT and *TMBIM5*^-/-^ HeLa cells in (A). Two-tailed *t*-test. A p-value of <0.05 was considered statistically significant. **(C)** Maximal respiration calculated from OCR experiment of WT and *TMBIM5*^-/-^ HeLa cells in (A). Two-tailed *t*-test. A p-value of <0.05 was considered statistically significant. **(D)** ATP production calculated from OCR experiments of WT and *TMBIM5*^-/-^ HeLa cells in (A). Two-tailed *t*-test. A p-value of <0.05 was considered statistically significant. **(E)** Mitochondrial ROS production. WT and *TMBIM5*^-/-^ HeLa cells were stained with MitoSox Red for 30 min and analyzed using fluorescence-activated cell sorting (FACS). Rotenone (0.5 µM) was added during MitoSox Red staining. Data are expressed as median (PE-A) ±SD (n=3 independent experiments). Two-tailed *t*-test. A p-value of <0.05 was considered statistically significant. **(F)** Enzymatic activities of different electron transport chain complexes (indicated as complex I = CI, complex II = CII, complex III = CIII and complex IV = CIV) measured in freeze thawed mitochondria from HEK293T WT and *TMBIM5*^-/-^ cells. Data are normalized to citrate synthase (CS) activities and expressed as mean ± SD percentage of control (n=3-4 independent mitochondrial preparations, each one measured in 3-6 technical replicates). Two-tailed *t*-test. A p-value of <0.05 was considered statistically significant. **(G)** Whisker-box plot showing log2 fold change distributions of proteins in HeLa *TMBIM5*^-/-^ and wildtype (WT) cells (n=5 independent experiments), which were identified in specific OXPHOS complexes I-IV. Boxes borders indicate the 25% and 75% quantiles and outliers are indicated by a + sign. AP= all cell proteome, M=MitoCarta 3.0, CI= complex I, CII= complex II, CIII= complex III, CIV= complex IV. **(H)** Bar graph displaying log2 fold changes between transcript (mRNA) and protein changes for subunits of the NADH dehydrogenase complex I. **(I)** Scatter plot comparing the turnover rate constants in in WT and *TMBIM5*^-/-^ HeLa cells. The incorporation rate was calculated and a first-order kinetic model was fitted to each peptide which allows the estimation of the turnover rate constant k [1/h]. For a protein group the mean peptide turnover constant was calculated. Subset of MitoCarta 3.0 assigned proteins are shown. Significantly different rate constants were identified using a two-sided t-test (p-value < 0.05). Complex I. subunits are highlighted in red. See also Table S4. **(J)** Cartoon of respiratory complex I highlighting the subunits affected by the deletion of *TMBIM5*. Orange subunits are part of N module (NDUFA2, NDUFA12, NDUFV2, NDUFS1). Green subunits are part of the Q module (NDUFA5, NDUFA9, NDUFS3, NDUFS7, NDUFS8 and the assembly factor NDUFAF4). The light blue subunit is part of the P module (NDUFB11). See also Figure S4 and Table S4.

To corroborate these findings, we analyzed the mitochondrial proteome in *TMBIM5*^-/-^ cells by mass spectrometry (Figure S4E, F, Table S2, 3). Deletion of *TMBIM5* was accompanied by significant alterations in the mitochondrial proteome (Figure S4E, F, Table S2, 3). STRING Gene Ontology enrichment analysis revealed that respiratory chain complexes, in particular complex I, were predominantly affected (Figure S4G). We therefore compared the steady state levels of all detected mitochondrial proteins and specifically those of subunits of the different respiratory complexes in wildtype and *TMBIM5*^-/-^ mitochondria (Figure 4G). Loss of TMBIM5 did not grossly alter the mitochondrial mass (Figure 4G) nor mtDNA levels (Figure S4H). However, subunits of respiratory complex I were significantly decreased in the absence of TMBIM5, in line with its reduced specific activity (Figure 4F). Subunits of other respiratory chain complexes were present at wildtype levels in *TMBIM5*^-/-^ mitochondria (Figure 4G). We therefore conclude that the loss of TMBIM5 has profound effects on the mitochondrial proteome and leads to reduced levels of respiratory complex I.

The decreased steady state level of complex I subunits in *TMBIM5* cells does not reflect reduced expression of these proteins (Figure 4H, Figure S4I), suggesting that TMBIM5 affects their stability. We therefore monitored the turnover of mitochondrial proteins upon stable isotope labeling with amino acids in cell culture (SILAC) (Ong et al., 2002) of wildtype and *TMBIM5*^-/-^ cells. After labeling cells with ^13^C- and ^15^N-labelled arginine and lysine, cells were further incubated in the presence of light isotopologues and the cellular proteome was determined at seven timepoints within 48 h by mass spectrometry (Figure S4J). We determined turnover rates of 348 mitochondrial proteins and identified 140 proteins, which were significantly faster turned over in *TMBIM5*^-/-^ cells relative to wildtype cells (Figure 4I, Table S4). The vast majority of these proteins are localized at the mitochondrial IM or matrix (131 out of 140). They include components of mitochondrial ribosomes and of the mitochondrial gene expression apparatus as well as various subunits of respiratory complex I, which were amongst the most significantly destabilized proteins (Figure 4I, Table S4).

We conclude from these experiments that TMBIM5 controls oxidative phosphorylation by affecting mitochondrial protein stability. Loss of TMBIM5 causes mitochondrial hyperpolarization, which is accompanied by an increased turnover especially of complex I (Figure 4J).

### TMBIM5 deletion promotes proteolysis by AFG3L2

Different turnover rates have been observed for the 44 subunits of complex I (Szczepanowska and Trifunovic, 2021). Subunits of the matrix-exposed N module are generally more rapidly degraded than membrane-bound core subunits and exchanged by newly synthesized subunits, likely to repair oxidative damage (Pryde et al., 2016; Szczepanowska et al., 2020). Degradation of N module subunits (NDUFS1, NDUFV1 and NDUFV2) is mediated by matrix-localized LON and CLPP proteases. The loss of TMBIM5 led to the proteolytic breakdown of 4 out of 5 identified subunits of the N-module (NDUFS1, NDUFA2, NDUFA12 and NDUFV2), but we also observed increased proteolysis of 6 out of 10 identified subunits of the Q module (NDUFA5, NDUFA9, NDUFS3, NDUFS7, NDUFS8 and the assembly factor NDUFAF4) as well as 1 out of 22 identified subunits of the P module in the IM (NDUFB11) (Figure 4I, J).

Since we have identified TMBIM5 as a binding partner of AFG3L2 (Figure 1A), we examined a possible involvement of AFG3L2 in the degradation of complex I subunits in TMBIM5-deficient mitochondria. An unbiased analysis of the proteolysis of OXPHOS is hampered by the known function of AFG3L2 for mitochondrial ribosome biogenesis and the synthesis of mitochondrially encoded respiratory chain subunits (Almajan et al., 2012). Rather than deleting *AFG3L2*, we therefore acutely depleted AFG3L2 from wildtype and *TMBIM5*^-/-^ mitochondria to limit effects on mitochondrial protein synthesis and determined the mitochondrial proteome by mass spectrometry. Among the 248 proteins whose steady state levels were decreased in mitochondria lacking TMBIM5 (Figures S4E), 215 proteins are localized to the IM or matrix space and therefore may include substrates of AFG3L2 (Figure 5A). Consistently, our unbiased proteomic approach revealed that steady state levels of 117 of the 215 proteins were significantly altered upon depletion of AFG3L2 in *TMBIM5*^-/-^ cells (Figure 5B, Table S5). Two protein classes can be distinguished: proteins that accumulate in the absence of AFG3L2 (cluster 1; 69 proteins) and proteins whose steady state levels further decrease upon downregulation of AFG3L2 (cluster 2; 48 proteins). 58 of the 69 cluster 1 proteins are localized to the matrix or IM and therefore represent candidate substrates of AFG3L2, whereas cluster 2 proteins are likely only indirectly affected by AFG3L2. Cluster 1 proteins also accumulate upon depletion of AFG3L2 in wildtype cells indicating AFG3L2-mediated turnover, which appears to be accelerated upon loss of TMBIM5 and mitochondrial hyperpolarization (Figure 5A).

**Figure 5.**
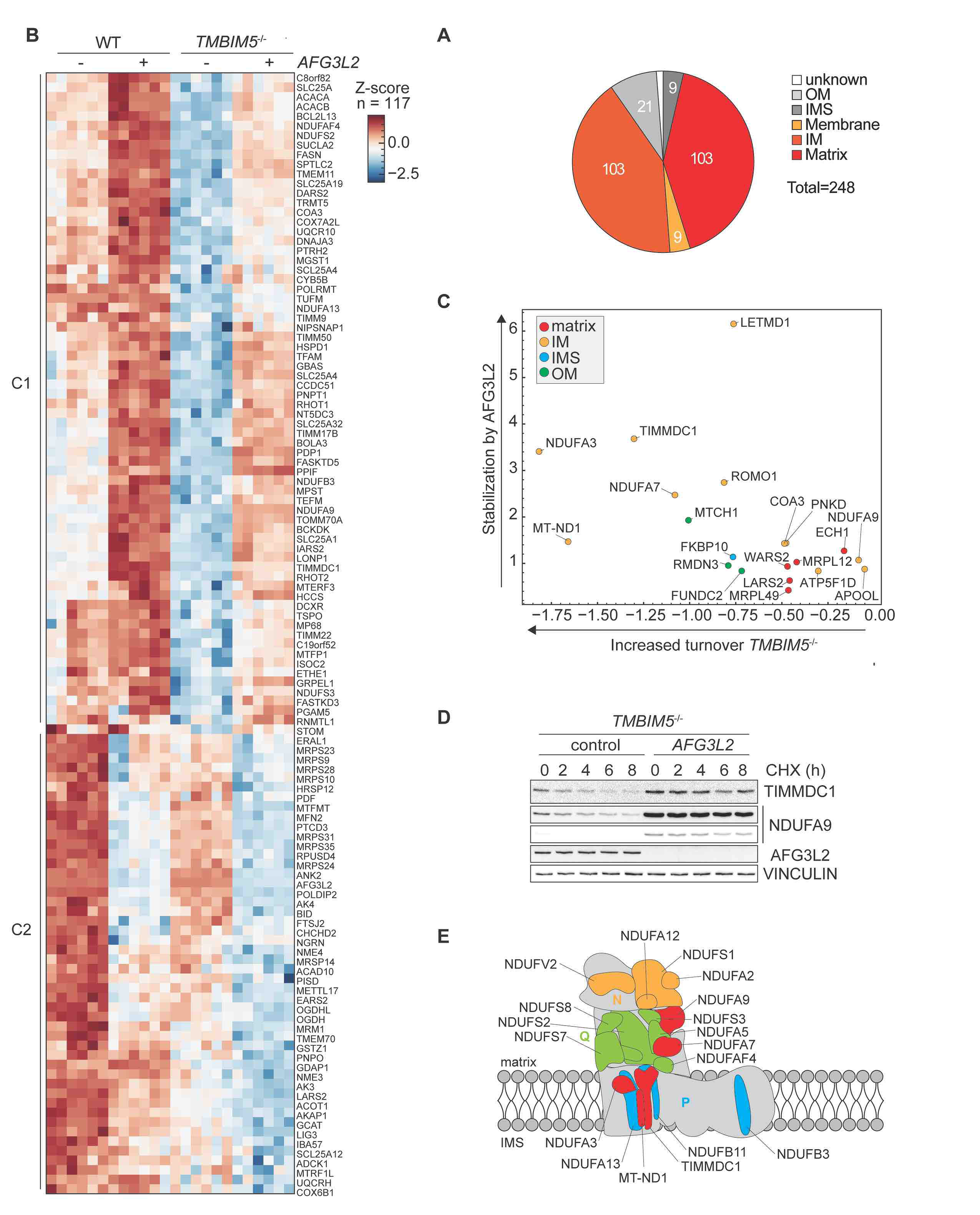
Loss of TMBIM5 promotes proteolysis by AFG3L2. **(A)** Pie chart showing the localization of proteins accumulating at decreased levels in *TMBIM5*^-/-^ HeLa cells. Localization according to MitoCarta 3.0. OM, outer membrane; IMS, intermembrane space; IM, inner membrane. See also Table S5. **(B)** Alterations in mitochondrial proteome in WT and *TMBIM5*^-/-^ HeLa cells upon depletion of AFG3L2. WT and *TMBIM5*^-/-^ HeLa cells were transfected with scrambled siRNA (control) or siRNA targeting *AFG3L2* for 48 h. Proteins, which were downregulated in the absence of TMBIM5 and whose steady state levels was altered upon depletion of AFG3L2, were identified using a two-sided t-test followed by permutation-based FDR estimation (0.05). Log2-transformed LFQ intensities of MitoCarta 3.0 proteins were Z-Score normalized and visualized in a hierarchical cluster analysis (Euclidean distance, complete method). The dendrogram is omitted. Proteins accumulating upon depletion of AFG3L2 (cluster 1, C1) and proteins whose steady state levels are decreased upon depletion of AFG3L2 (cluster 2, C2) are shown. See also Table S5. **(C)** Scatter plot showing the area under curve (AUC) difference between cycloheximide chase protein abundance data between (WT, scrambled) and (*TMBIM5*^-/-^, scrambled) rate in *TMBIM5*^-/-^ HeLa cells (x-axis) versus the AUC difference between (*TMBIM5*^-/-^ control) and (*TMBIM5*^-/-^ siAFG3L2). Only MitoCarta 3.0 proteins are displayed. See also methods for data analysis and Table S6. **(D)** Representative immunoblot of *TMBIM5*^-/-^ HeLa cells transfected with scrambled siRNA (control) or siRNA targeting *AFG3L2* for 48 h. Samples were treated with the protein synthesis inhibitor cycloheximide (CHX; 10 μg/ml) and collected at the indicated time points (n=3 independent experiments). **(E)** Cartoon of respiratory complex I highlighting subunits affected by *TMBIM5* deletion. Subunits of the N module (NDUFV2, NDUFA12, NDUFS1, NDUFA2) are shown in orange, subunits of the Q module (NDUFS2, NDUFS3, NDUFA5, NDUFS8, NDUFS7 and the assembly factor NDUFAF4) in green and subunits of the P module (NDUFB11, NDUFB3, NDUFA13) in blue. Red color is used for complex I subunits that are AFG3L2 substrates according to (C) (NDUFA3, NDUFA7, NDUFA9, MT-ND1 and TIMMDC1). See also Table S5 and S6.

To directly assess the effect of TMBIM5 on AFG3L2-mediated proteolysis, we examined how downregulation of AFG3L2 affected the stability of mitochondrial proteins in *TMBIM5*^-/-^ cells. Since the strong growth phenotype of AFG3L2-deficient cells precluded pulse-SILAC experiments, we inhibited cytosolic protein synthesis with cycloheximide in wildtype and *TMBIM5*^-/-^ cells and analyzed the alterations in the mitochondrial proteome by mass spectrometry after 0, 2, 4, 6 and 8 h treatment in control or AFG3L2 knockdown conditions using a data independent acquisition approach in 6 biological replicates. In total, we identified 10,150 protein groups out of which 990 protein groups are localized to mitochondria (MitoCarta 3.0) indicating a comprehensive coverage of the proteome. After calculating relative protein abundance over time, we determined the area under fitted curves, which indicates protein stability and is increased if proteolysis is accelerated. Using stringent selection criteria, this approach identified 20 proteins, which were faster degraded in *TMBIM5*^-/-^ cells but stabilized upon downregulation of AFG3L2 in a statistically significant manner (Figure 5C, Table S6). IM, matrix and OXPHOS related proteins were enriched among the stabilized proteins, including complex I subunits NDUFA3, NDUFA7, NDUFA9, MT-ND1 and the complex I assembly factor TIMMDC1 that was recently identified as AFG3L2 substrate (Wang et al., 2021)(Figure 5C, Table S6). Immunoblot analysis of cellular fractions after inhibition of cytosolic protein synthesis confirmed the AFG3L2-dependent degradation of TIMMDC1 and NDUFA9 (Figure 5D).

We conclude from these experiments that the loss of TMBIM5 causes broad reshaping of the mitochondrial proteome and promotes the AFG3L2-mediated degradation of IM and matrix proteins, in particular subunits of respiratory complex I (Figure 5E).

### Hyperpolarization induces TMBIM5 degradation

It is conceivable that mitochondrial hyperpolarization in the absence of TMBIM5 triggers proteolysis. To examine this possibility, we induced persistent mitochondrial hyperpolarization by inhibiting the F1FO ATP synthase with oligomycin in wildtype cells and monitored effects on the mitochondrial proteome by mass spectrometry (Figure 6A, B; Figure S5A: Table S7). Surprisingly, only 87 mitochondrial proteins were significantly downregulated upon oligomycin treatment, i.e. significantly less proteins than in *TMBIM5*^-/-^ cells (Figure 6A, Table S7). Only 35 of the proteins were affected by both oligomycin treatment and deletion of *TMBIM5* (Figure 6A, Table S7). Thus, mitochondrial hyperpolarization is not sufficient to explain the broad effect of TMBIM5 on the mitochondrial proteome.

**Figure 6.**
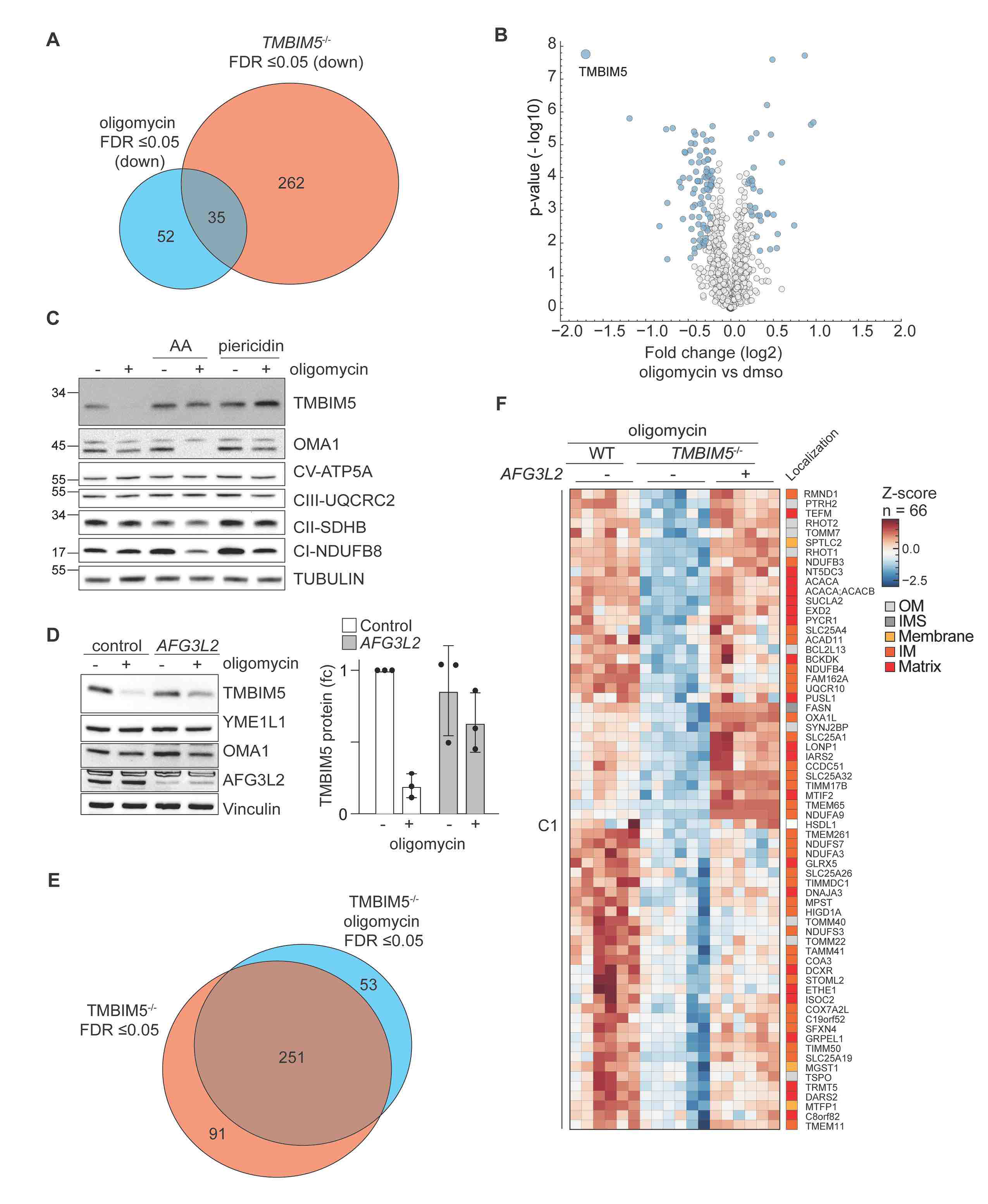
Hyperpolarization induces degradation of TMBIM5 allowing AFG3L2- mediated remodeling of the mitochondrial proteome. (A) Venn diagram of significantly downregulated proteins (FDR *≤*0.05) in oligomycin-treated WT HeLa cells (0.01% DMSO vs oligomycin 10 μM for 16 h; blue) and in *TMBIM5*^-/-^ HeLa cells (WT vs *TMBIM5*^-/-^ HeLa cells; red). See also Table S7. (B) Volcano plot of mitochondrial protein (MitoCarta 3.0) changes in WT HeLa cells treated with oligomycin (10 μM for 16 h) when compared to untreated cells. Before MitoCarta 3.0-filtering, significantly altered proteins were identified using a two-sided t-test followed by permutation-based FDR estimation to 0.05 and are colored in blue (n=5 independent experiments). See also Table S7. (C) Representative immunoblot of WT HeLa cells treated with the indicated drugs for 16 h. Antimycin A (AA; 10 μM), piericidin A (10 μM) and oligomycin (10 μM). Quantification is shown in Figure S5B (n=7 independent experiments). (D) Representative immunoblot of WT HeLa cells transfected with scrambled siRNA (control) and siRNA targeting *AFG3L2* for 48 h, treated when indicated with oligomycin (10 μM) for 16 h. Quantification of TMBIM5 levels is shown on the right panel. TMBIM5 levels in untreated WT cells was set to 1 (n=3 independent experiments). (E) Venn diagram of significantly downregulated proteins (FDR *≤*0.05) in oligomycin-treated *TMBIM5^-/-^* HeLa cells (0.01% DMSO vs oligomycin 10 μM for 16 h; blue) and in HeLa cells (WT vs *TMBIM5*^-/-^ HeLa cells; red). See also Table S5. (F) Zoom in to selected clusters of hierarchical clustering of Z-Score normalized log2 LFQ intensities. Proteins that were significantly (FDR < 0.05) accumulated between HeLa WT and TMBIM5^-/-^ cells after oligomycin treatment (10µM) for 16 h were clustered using Euclidean distance and the complete method on protein features. The clustering is shown in Suppl Table S5. The figure shows identified cluster C4 and C5 with gene names and MitoCarta 3.0 localization annotations. The row dendrogram is omitted. See also Table S5. See also Figure S6 and Tables S5 and S7.

We noticed that among all detected mitochondrial proteins TMBIM5 was the strongest decreased protein upon mitochondrial hyperpolarization (Figure 6B, Table S7), although TMBIM5 transcript levels were not significantly altered under these conditions (Figure S5B- D). We reasoned that pH-dependent conformational changes in the presence of oligomycin may destabilize TMBIM5, as it has been suggested for other TMBIM family members (Saraiva et al., 2013). We therefore incubated oligomycin-treated cells additionally with other OXPHOS inhibitors, which limit H^+^ transport and thus prevent matrix alkalinization and hyperpolarization of the IM (Figure 6C). Inhibition of respiratory complex I with piericidin A or of respiratory complex III with antimycin A indeed stabilized TMBIM5 in cells treated with oligomycin but did not affect TMBIM5 levels in untreated cells (Figure 6C, Figure S5B-D). These results reveal that hyperpolarization triggers proteolysis of TMBIM5, which is abolished upon depolarization of mitochondria. Degradation of TMBIM5 in hyperpolarized mitochondria is at least in part mediated by AFG3L2. Downregulation of AFG3L2 slowed degradation and stabilized TMBIM5 in the presence of oligomycin (Figures 6D; Figure S5E). Notably, AFG3L2 depletion did not depolarize mitochondria under these conditions, excluding indirect effects on TMBIM5 stability (Figure S5F). We conclude that TMBIM5 is degraded by AFG3L2 upon persistent hyperpolarization of mitochondria.

### Hyperpolarization and TMBIM5 degradation allow remodeling of the mitochondrial proteome by AFG3L2

These experiments identify TMBIM5 both as an interactor and substrate of AFG3L2, which mediates TMBIM5 turnover upon mitochondrial hyperpolarization. This is reminiscent of *E*.

*coli* YccA, a distant homologue of the TMBIM family, which shares 12.5% sequence identity with TMBIM5. YccA is degraded by FtsH, the bacterial homologue of AFG3L2, while its binding to FtsH inhibits proteolysis of other FtsH substrates (Kihara et al., 1998; van Stelten et al., 2009). Assuming functional conservation, we hypothesized that TMBIM5 binding inhibits AFG3L2 activity, while its degradation in hyperpolarized mitochondria allows broad reshaping of the mitochondrial proteome by AFG3L2 mediated proteolysis. The degradation of TMBIM5, acting as a competitive inhibitor, may restrict AFG3L2 activity upon acute hyperpolarization. However, progressive TMBIM5 degradation upon persistent hyperpolarization is expected to increasingly release AFG3L2 activity towards other substrates and drive the proteolytic breakdown of mitochondrial proteins.

To examine this hypothesis, we first assessed how oligomycin-induced hyperpolarization affects the mitochondrial proteome in *TMBIM5*^-/-^ cells. In contrast to our previous observations in wildtype cells (Figure 6A), oligomycin broadly altered the proteome of *TMBIM5*^-/-^ cells, showing decreased levels of 304 proteins (Figure 6E). Strikingly, the levels of a large fraction of them (251 proteins) were reduced in untreated *TMBIM5*^-/-^ cells (Figure 6E, Table S5). These experiments suggest that the presence of TMBIM5 limits oligomycin-induced changes in the mitochondrial proteome of wildtype cells (Figure 6A), consistent with TMBIM5 inhibiting AFG3L2. We therefore analyzed in further experiments how depletion of AFG3L2 affected mitochondrial proteins in oligomycin-treated *TMBIM5*^-/-^ cells (Figure 6F). We identified 66 proteins, whose steady state levels were significantly decreased in hyperpolarized *TMBIM5*^-/-^ cells when compared to wildtype cells, but which accumulated upon depletion of AFG3L2 (Figure 7B, Table S5). 56 of these proteins, including 8 subunits and assembly factors of respiratory complex I (NDUFA3, NDUFA9, NDUFS3, NDUFS7, NDUFB3, NDUFB4, TMEM261 and TIMMDC1), are localized in the IM or matrix of mitochondria and therefore likely represent AFG3L2 substrates in hyperpolarized mitochondria. These data demonstrate increased proteolysis by AFG3L2 in hyperpolarized mitochondria lacking TMBIM5 substantiating the regulation of AFG3L2 by TMBIM5 and the proton gradient.

**Figure 7.**
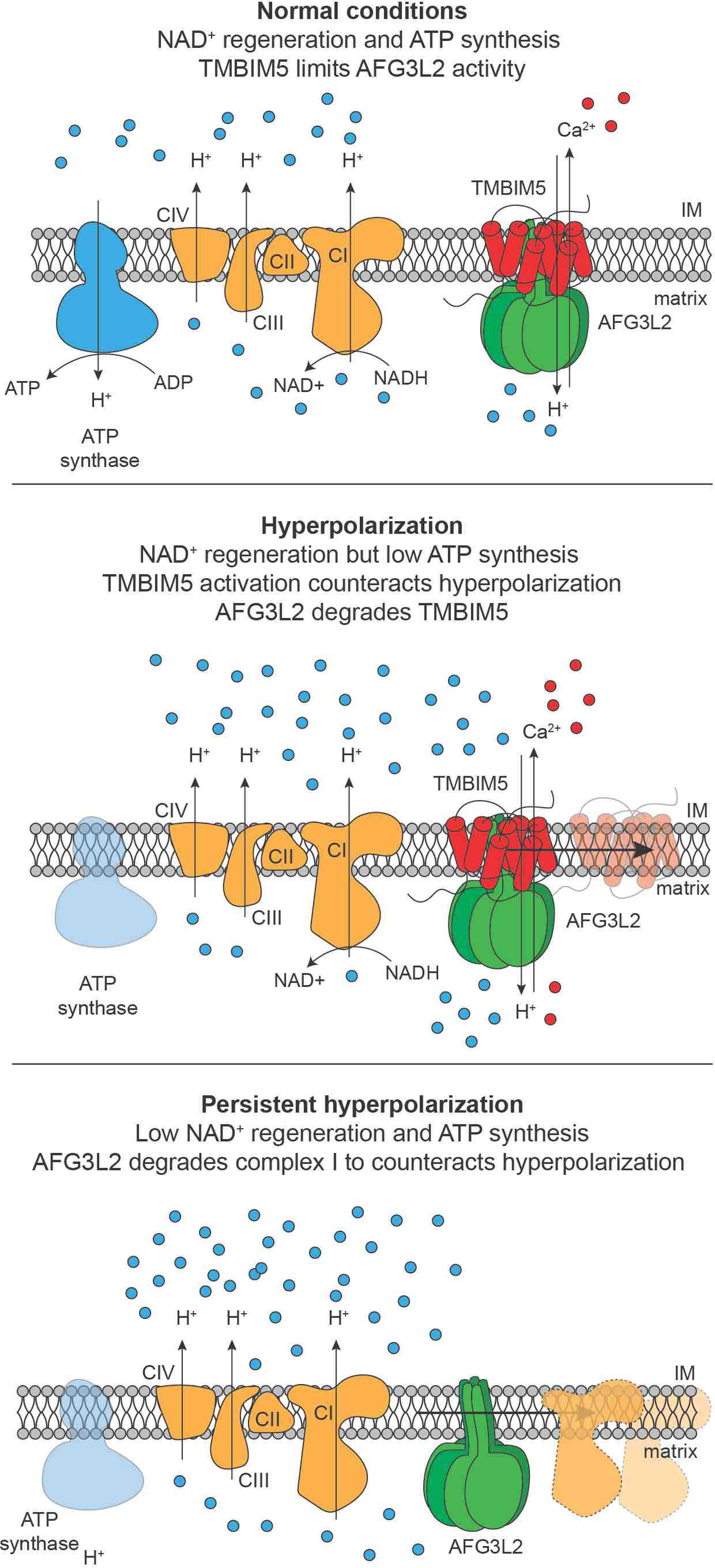
Model for the regulation of AFG3L2-mediated proteolysis by TMBIM5 under different metabolic conditions. Upper panel, TMBIM5 inhibits AFG3L2 and, acting as a Ca^2+^/H^+^ exchanger, limits the formation of the mitochondrial H^+^ gradient. Middle panel, TMBIM5 counteracts acute hyperpolarization and limits ROS production, for instance, if the demand for mitochondrial ATP is low. Moreover, TMBIM5 inhibits AFG3L2-mediated proteolysis allowing electron flow through complex I and NAD^+^ regeneration. Lower panel, persistent mitochondrial hyperpolarization induces the proteolytic breakdown of TMBIM5 and releases AFG3L2 activity, which reshapes the mitochondrial proteome and degrades respiratory complex I subunits to prevent excessive ROS production.

## Discussion

We have identified TMBIM5 as a mitochondrial Ca^2+^/H^+^ exchanger, which binds to and inhibits the m-AAA protease AFG3L2 and therefore couples mitochondrial protein turnover with the energetic status of mitochondria (Figure 7). TMBIM5 allows Ca^2+^ efflux from mitochondria counteracting Ca^2+^ overload and apoptosis. Moreover, TMBIM5 activity limits the formation of the H^+^ gradient across the IM and inhibits proteolysis by the m-AAA protease. Persistent mitochondrial hyperpolarization induces the degradation of TMBIM5 and activation of the m-AAA protease associated with broad remodeling of the mitochondrial proteome. The resulting degradation of complex I subunits limits ROS production in hyperpolarized mitochondria and contributes to the rebalancing of the H^+^ gradient.

We used cellular and *in vitro* reconstitution experiments to demonstrate that TMBIM5 mediates Ca^2+^ efflux across the IM, which is accompanied by H^+^ transport into the mitochondrial matrix. Thus, TMBIM5 appears to exert Ca^2+^/H^+^ exchange activity, while other members of the TMBIM family act as pH-dependent Ca^2+^ channels in membranes lacking a stable H^+^ gradient (Bultynck *et al*., 2014; Guo *et al*., 2019; Li *et al*., 2020). Similar to the Na^+^/Ca^2+^ exchanger NCLX (Katoshevski et al., 2021; Palty et al., 2010), TMBIM5 allows the efflux of Ca^2+^ from mitochondria, raising the question why different Ca^2+^ efflux routes have evolved. Although TMBIM5 is ubiquitously expressed in mice, tissue-specific demands may exist. Moreover, both TMBIM5 and NCLX affect OXPHOS activity, but exchange Ca^2+^ against different cations. Na^+^ influx through NCLX directly modulates complex II and III activity by affecting membrane fluidity (Hernansanz-Agustin et al., 2020). We demonstrate here that TMBIM5 preserves respiration regulating the formation of the H^+^ gradient across the IM. Accordingly, loss of TMBIM5 leads to mitochondrial hyperpolarization and impairs OXPHOS activities. We propose that coupling of cycles of Ca^2+^ influx via MCU and Ca^2+^ efflux via TMBIM5 allows tuning the H^+^ gradient across the IM independent of the F1FO-ATP synthase. This pathway may be of relevance when the demand for NAD^+^ exceeds that for mitochondrial ATP, as observed during aerobic glycolysis (Luengo et al., 2021). Cell growth does not depend on mitochondrial ATP in glycolytic cells, but NAD^+^ regeneration at respiratory complex I maintains the cellular redox homeostasis and the activity of numerous NAD^+^ dependent enzymes (Covarrubias et al., 2021). Under conditions of limited mitochondrial ATP production, dissipation of the H^+^ gradient by TMBIM5 counteracts hyperpolarization (Figure 7). TMBIM5 thereby preserves H^+^ transport by respiratory complexes and facilitates NAD^+^ regeneration at complex I, while at the same time limiting ROS production. The Ca^2+^/H^+^ exchange activity of TMBIM5 therefore allows to utilize the proton gradient for NAD^+^ regeneration independent of the F1FO-ATP synthase.

This metabolic role of TMBIM5 is supported by its second activity as inhibitor of AFG3L2. Inhibition of the m-AAA protease prevents the premature degradation of unassembled or damaged respiratory chain subunits and therefore maintains respiration. We identify TMBIM5 as a substrate of AFG3L2, whose proteolysis inhibits the degradation of other AFG3L2 substrates upon acute hyperpolarization. Only persistent hyperpolarization leads to the complete loss of TMBIM5 and the release of m-AAA protease activity (Figure 7). It is conceivable that hyperpolarization induces conformational changes in TMBIM5, reminiscent of other TMBIM family members. Physiological pH changes were shown to trigger TMBIM4 oligomerization regulating its anti-apoptotic function (Saraiva *et al*., 2013). In the absence of TMBIM5, the m-AAA protease broadly reshapes the mitochondrial proteome. Our unbiased proteomic approaches identify >100 proteins that are degraded in an AFG3L2-dependent manner, highlighting the central role of the m-AAA protease for mitochondrial proteostasis. Degradation of subunits of complex I, which are among the most affected proteins in the absence of TMBIM5, limits respiration and ROS production. Previous studies have already revealed rapid degradation of N-module subunits of complex I upon mitochondrial damage and established the involvement of the matrix-localized LON and CLPP peptidases (Pryde *et al*., 2016; Szczepanowska *et al*., 2020). Our experiments are consistent with and complement these findings and demonstrate a key role of the m-AAA protease in the degradation of complex I. Thus, various mitochondrial quality control peptidases are involved in the turnover of complex I subunits. It is an attractive possibility that the topology of the subunits within complex I and the stability of sub-complexes determine the involvement of a protease in their proteolytic breakdown. Consistently, increased AFG3L2-mediated proteolysis does not solely explain the broad alterations in the mitochondrial proteome in the absence of TMBIM5. Other mitochondrial peptidases might be activated upon mitochondrial hyperpolarization, although indirect effects of the loss of TMBIM5 and increased AFG3L2-mediated proteolysis cannot be excluded.

Similar to the regulation of the m-AAA protease by TMBIM5, *E. coli* YccA distantly related to TMBIM5 inhibits the bacterial homologue of the m-AAA protease FtsH (Kihara *et al*., 1998; van Stelten *et al*., 2009). Although an ion transport activity of YccA has not been demonstrated, YccA shares limited sequence identity with the pH-sensitive domain of TMBIM proteins, suggesting conserved regulation of FtsH-like proteases from bacteria to human. Notably, a homologue of mitochondrial TMBIM5 is not present in the yeast *S. cerevisiae*, which also lacks MCU as the major Ca^2+^ uptake route into mitochondria as well as respiratory complex I, pointing to a co-evolution of the Ca^2+^ dependent regulation of proteolytic activity with these complexes. Regardless, our results identify the m-AAA protease as a central hub shaping Ca^2+^ cycling into mitochondria. The m-AAA protease limits Ca^2+^ efflux from hyperpolarized mitochondria by degrading TMBIM5 and, at the same time, prevents mitochondrial Ca^2+^ overload by the proteolytic breakdown of excess EMRE subunits which ensures the formation of gated MCU-EMRE channels only (Konig *et al*., 2016; Tsai *et al*., 2017). It will be of interest to define the role of these regulatory circuits in neurodegenerative disorders associated with mutations in m-AAA protease subunits. Considering the prominent role of mitochondrial Ca^2+^ homeostasis for mitochondrial metabolism, cell signaling and cell death pathways (Garbincius and Elrod, 2021), TMBIM5 acting as a Ca^2+^/H^+^ exchanger and inhibitor of the m-AAA protease is likely of central importance both in healthy and pathophysiological conditions.

## Author contributions

M.P. and T.L. wrote the manuscript; M.P. designed and performed most of the experiments and analyzed the data with contributions from Y.L., T.K. and A.E.; D. T. prepared liposomes containing TMBIM5; M. G. and M.M. performed electrophysiology and analyzed the data;H.N. and M.P. designed, performed and analyzed the proteomic experiments; Y.O. generated TMBIM5 HeLa KO cells; Y.L. performed AFG3L2 interactome; Z.A.A., A.C-O and U.B. performed the analysis of respiratory chain complex activities. All authors commented on and edited the manuscript.

## Acknowledgments

We thank Kathrin Lemke and Dominique Diehl for excellent technical assistance. Support in the EM analysis by Janine Heise (CECAD imaging facility) and in the FACS analysis by Kat Folz-Donahue (MPI imaging facility) is greatly appreciated. This work was supported by an EMBO postdoctoral fellowship to M.P. (ALTF 649-2015; LTFCOFUND2013; and GA-2013- 609409), a fellowship of the Japan Society for the Promotion of Science (JSPS) for research abroad, The Osamu Hayaishi Memorial Scholarship for Study Abroad and grants from the Uehara Memorial Foundation to Y.O., grants by the European Union’s Horizon 2020 research and innovation programme under the Marie Sklodowska-Curie grant agreement No 721757 and the German-Israel-Project (DIP, RA1028/10-2) to T.L., grants from the Deutsche Forschungsgemeinschaft (FOR2848; projects A01 and A05) to T.L. and M.M., respectively, and grants from the Netherlands Organization for Scientific Research (TOP 714.017.00 4) and from the Deutsche Forschungsgemeinschaft (CRC1218 - Project number 269925409) to U.B..

## Material and Methods

### Cell culture

If not otherwise indicated HeLa, HEK 293T and Flp-In HEK293 T-REx were maintained in DMEM-GlutaMAX at 4.5 g*ml^-1^ D-glucose (Gibco) supplied with sodium-pyruvate, non-essential amino acids (both Gibco) and 10% fetal bovine serum (Sigma). Cells were seeded with equal densities before an experiment and regularly checked for Mycoplasma contamination.

## Method details

### Reagents

Oligomycin (Cat# O4876), Antimycin A (Cat# A8674), nigericin (Cat# N7143), cycloheximide (Cat# C4859), rotenone (Cat# R8875), ionomycin (Cat# 13909), thapsigargin (Cat# T9033), staurosporine (Cat# S6942), Actinomycin D (Cat# A9415) were purchased from Sigma Aldrich. Piercidin (Cat# SC-202287) was purchase from Chemcruz. Venetoclax (Cat# S8048) and A-1155463 (S7800) were purchased from Selleckchem. Fura2 (Cat# F1200), TMRM (Cat# T668) and MitoSox (Cat# M36008) were purchased from Invitrogen. Calcium-Green-5N (Cat# C3737) was purchased from life technologies.

### Antibodies

The following antibodies were used for immunoblotting: PARP (ThermoFisher, Cat# PA-951; 1:1000), TMBIM5 (Proteintech, Cat# 16296-1-AP; 1:2000), VINCULIN (Cell signaling, Cat# 4650; 1:1000), MCU (Sigma, Cat# HPA016480; 1:1000), PHB1 (NeoMarkers, Cat# RB-292; 1:500), AFG3L2 (Sigma, Cat# HPA004480; 1:5000), ATPVa (Abcam, Cat# Ab14748; 1:5000), TIMMDC1 (Abcam, Cat# Ab171978; 1:1000), NDUFA9 (Invitrogen, Cat# 459100, 1:1000), OMA1 (Santa Cruz, Cat# SC-515788; 1:1000), Total oxphos rodent (Abcam, Cat# Ab110413, 1:40000), YME1L1 (Proteintech, Cat# 11510-1-AP; 1:2000), Tubulin (Sigma, Cat# T6074; 1:5000). The following antibodies were used for immunofluorescence: cytochrome c (ThermoFisher, 33-8200; 1:500) and TOMM20 (Sigma, Cat# WH0009804M1; 1:500). Corresponding species-specific HRP-coupled antibodies were used for immunoblot (Biorad; Cat# 1706515 and Cat# 1706516) and fluorescently coupled secondary antibodies were used for immunofluorescence (Invitrogen Alexa Flour).

### DNA plasmids

Complementary human cDNA encoding human TMBIM5 and AFG3L2-FLAG was cloned into pcDNA5/FRT/TO (Invitrogen). AFG3L2 was mutated to AFG3L2(E408Q) and TMBIM5 to TMBIM5 D294L D325K via site-directed mutagenesis.

### Generation of stable expression cell lines

Flp-In HEK293T-REx cells were transfected with pcDNA5/FRT/TO AFG3L2^E408^FLAG. After transfection cells were selected with 100 ug/ml hygromycin (Roth, Cat# CP13.4) for 7 days. Single colonies were selected and expression of the gene of interest was induced with 1 mg/ml tetracycline (Roth, Cat# 0237.1) for 24 h before testing expression via immunoblotting.

### CRISPR-Cas9 gene editing

For generation of *TMBIM5*^-/-^ cell lines HeLa cells were used. Cells were transfected with px335 (Addgene #42335) plasmids containing either (3’-CCTCAGTATGTCAAGGATAGAAT-5’) or (3’-CCACCTATATGTACTTAGCAGGG-5’) guide RNA for *TMBIM5*. After three transfections cells were seeded as single cells in 96-well plates and checked for TMBIM5 protein expression after three weeks. Additionally, the *TMBIM5* mRNA level was tested via quantitative real-time PCR and the genomic region of *TMBIM5* was sequenced for validation of the knock out. Gene editing results in three different mutations in exon V and insertion of a premature STOP codon.

### Plasmid and siRNA transfection

For transient expression of DNA, GeneJuice transfection reagent (Merck) was used. Experiments were conducted 24 h after transfection. The following siRNA targeting sequences against human proteins were purchased from (Merck): TMBIM5 (SASI_HS01_00196210), MCU (SASI_HS01_00234629), AFG3L2 (SASI_HS01_0023-1520), NCLX (SASI_HS01_00236391). Negative control medium GC duplex was purchased from (Invitrogen; Cat# 12935300). RNA interference experiments with siRNA were conducted with Lipofectamine RNAiMAX (Thermofisher) transfection reagent. Efficient protein depletion was tested either by immunoblotting or by RT-PCR. Results shown in the paper related to depletion of TMBIM5 by siRNA represent the results obtained using one siRNA, which was selected among 5 different siRNA sequences, all of them showing similar effects. Cells were grown for either 48 or 72 h after transfection, as indicated in the Figure legends.

### Measurement of cellular respiration

Oxygen consumption rate was measured in a Seahorse Extracellular Flux Analyzer XFe96 (Agilent) in cells grown in DMEM-GlutaMAX containing 25 mM glucose. 4*10^4^ cells were plated in each well and incubated 1 h before measurement at 37°C. ATP production was assessed after addition oligomycin (2 µM); maximal respiration after FCCP (0.5 µM) and non-mitochondrial respiration after addition of rotenone and antimycin A (0.5 µM).

### Activities of respiratory chain complexes

HEK293T WT and *TMBIM5*^-/-^ cells were cultured in DMEM (4.5 g/L glucose; Lonza) supplemented with 10% (v/v) fetal bovine serum, 1% (v/v) penicillin (10,000 units/ml)/ streptomycin (10 mg/ml), 1 mM sodium pyruvate and 1% non-essential amino acids at 37 °C, 5% CO2. Confluent cells were harvested and resuspended in ice-cold isolation buffer (250 mM sucrose, 0.1% bovine serum albumin (BSA), 1 mM EDTA, 20 mM Tris-HCl, pH 7.4) and disrupted by ten strokes using a Potter-Elvehjem homogenizer with a Teflon pestle. After a clarifying spin at 1,000 x *g* for 10 min at 4°C, crude mitochondria were harvested from supernatant by sedimentation at 10,000 x *g* for 10 min at 4°C. For further purification, the mitochondrial pellet was resuspended in isolation buffer and loaded on a two-layer (1 M and 1.5 M) sucrose gradient buffered with 1 mM EDTA, 20 mM Tris/HCl, pH 7.4. After centrifugation for 20 min at 60,000 x *g* (4°C), the mitochondrial layer localized at the top of the 1.5 M sucrose cushion was carefully collected and washed twice with BSA-free isolation buffer by centrifugation at 22,000 x *g* (10 min; 4°C). Finally, mitochondria were resuspended in a small volume of BSA-free isolation buffer. Protein concentration was determined by the Lowry method and mitochondria were shock-frozen in liquid N2 and stored at -80°C until use.

For the kinetic measurements, mitochondria were diluted to 1 mg/ml in 20 mM Tris-HCl, pH 7.2 and subjected to three freeze-thaw cycles to break the inner membrane. Activities of the individual respiratory chain complexes and of citrate synthase as a reference were measured spectrophotometrically at 25°C in a SPECTRAmax PLUS^384^ microplate reader (Molecular Devices). Complex I (NADH:*n*-decylubiquinone (DBQ) oxidoreductase) activity: 20 µg/ml mitochondria, 200 µM NADH, 70 µM DBQ, 3 mg/ml BSA, 2 µM antimycin A, 500 µM NaCN, 25 mM potassium phosphate, pH 7.5. NADH oxidation was followed at 340 nm (*ε* = 6.2 mM^-1^·cm^-1^) and corrected for the inhibitor-insensitive rate determined in the presence of 10 µM 2-decyl-4-quinazolinyl amine (DQA). Complex II (succinate:2,6-dichlorophenol- indophenol (DCPIP) oxidoreductase) activity: 20 µg/ml mitochondria, 10 mM Succinate, 70 µM DBQ, 80 µM DCPIP, 1 mg/ml BSA, 2 µM myxothiazol, 500 µM NaCN, 20 mM Tris-HCl pH 7.2. DCPIP reduction was followed at 600 nm (*ε* = 19.1 mM^-1^·cm^-1^) and corrected for the inhibitor-insensitive rate determined in the presence of 10 mM malonate. Complex III (ubiquinol:cytochrome *c* oxidoreductase) activity: 20 µg/ml mitochondria, 75 µM ferricytochrome *c*, 100 µM *n*-decylubiquinol (DBQH2), 0.1 mM EDTA, 500 µM NaCN, 25 mM potassium phosphate, pH 7.5. Cytochrome *c* reduction was followed at 550 nm (*ε* = 18.5 mM^-1^·cm^-1^) and corrected for the inhibitor-insensitive rate determined in the presence of 2 µM myxothiazol. Complex IV (cytochrome *c* oxidase) activity: 20 µg/ml mitochondria, 50 µM ferrocytochrome *c*, 25 mM potassium phosphate, pH 7.0. Cytochrome *c* oxidation was followed at 550 nm (*ε* = 18.5 mM^-1^·cm^-1^) and corrected for the inhibitor-insensitive rate determined in the presence of 500 µM NaCN. Citrate synthase activity: 10-20 µg/ml mitochondria, 1 mM oxaloacetate, 0.3 mM Acetyl-CoA, 0.1 mM 5’-dithiobis 2-nitrobenzoic acid (DTNB), 0.1% Triton X-100, 100 mM Tris-HCl, pH 8.0. The reaction of DTNB with CoA formed upon citrate formation was followed at 412 nm (*ε* = 13.6 mM^-1^cm^-1^).

### Immunofluorescence

HeLa cells transfected with indicated constructs were plated on 13 mm glass coverslips in 24 well dishes. Cells were washed with room temperature PBS and fixed with 3.7% PFA in PBS at room temperature for 20 min. Cells were permeabilized in PBS+0.2% Triton X-100. Samples were blocked in PBS+0.1% BSA for 30 min before incubating over night with primary IF- antibody reconstituted in blocking solution with dilutions indicated above. The secondary IF- antibody (Invitrogen, Alexa Flour) was reconstituted in blocking buffer and incubated for 1 h, before washing the samples and mounting the coverslips onto microscopy slides with Prolong Gold Antifade Mountant (Invitrogen, Cat# P10144). Image acquisition was performed with a Leica SP8-DLS laser-scanning confocal microscope using an 100x oil objective (Leica HC PL APO 100×/1.4 OIL CS2; Cat# 11506372).

### pH and Ca*^2+^* measurements

Cells were plated on glass coverslips and transfected either with SypHer3s-dmito (Addgene #108119) or with mtRED-GECO (Addgene #46021). Imaging with a Leica SP8- DLS laser-scanning confocal microscope using an 40x oil objective (Leica HC PL APO 40×/1.3 OIL) was performed 24 h after transfection with the following protocol. 24 h after transfection coverslips were mounted into custom made open topped-chamber and medium exchanged with freshly prepared KRB buffer (135 mM NaCl, 5 mM KCl, 1 mM MgCl2, 20 mM HEPES, 1 mM MgSO4, 0.4 mM KH2PO4, 1 mM CaCl2, 5.5 mM glucose; pH 7.4 using NaOH) if not otherwise indicated at 37C. Sypher was excited at a wavelength of 405 nm and 488 nm, mtRED-GECO at a wavelength of 550 nm. Calibration of SypHer3s-dmito was performed in calibration buffer (130 mM KCl, 10 mM NaCl, 2 mM K2HPO4, 1mM MgCl2) supplemented with 20 mM MES buffer (adjusted to pH 5.5 and 6.5 with KOH) or 20 mM HEPES buffer (adjusted to pH 7.0 and 7.5 with KOH) or TRIS buffer (adjusted to pH 8.0 and 9.0 with HCl) or Acid Boric (adjusted to pH 9.5 and 10 with KOH), pH was stepped between and 10 by turnover of the bath solution. For each experiment monensin (5 μM) and nigericin (1 μM) were also added. The ratio of absorbance at 488 nm and 405 nm was calculated. Images were processed in FiJi. For each cell, a 8-point calibration curve was fitted to a variable slope sigmoid equation.

Ca^2+^ experiments were also performed in KRB buffer. Medium was changed with KRB 10 min before the experiment. Images were acquired every 1.5 s. Thapsigargin (2 μM) was added after 200 s. Data are show as fluorescence (F) F/F0 quantification, where F0 is the average of the first 10 sec and F is the fluorescence peak measured after thapsigargin administration.

### Mitochondrial membrane potential

5*10^5^ cells were incubated with TMRM (Thermofisher; 200 nM) containing CCCP (15 μM) or DMSO (0.01%) for 30 min at 37°C. Cells were washed with PBS, transferred in a black 96 well plate and the fluorescence was analyzed with a plate reader (Perkin Elmer EnVision 2105).

### Quantitative Real-Time PCR

Isolation of mRNA from HeLa cells was performed with NucleoSpin RNA kit (Macherey-Nagel; 740955.250) before generating cDNA with Promega GoScript Reverse Transcriptase kit (Cat# A5003) and adjusting to 10 µg/ml. PCR was performed with SYBR Green PCR mix (applied biosystems; Cat# 4309155), 0.6 µM of each primer as well as 30 ng cDNA for 40 rounds at 95°C melting and 60°C elongation steps. Actin and GADPH was used as housekeeping genes for normalization. Primer sequences are listed in the key resource table.

### SDS-PAGE and immunoblotting

Cells were collected and washed with PBS, before lysing with lysis-buffer (1 M (w/v) Tris pH 8.5, 0,5 M (w/v) EDTA, 10% (w/v) SDS, 3 M (w/v) NaCl) containing complete protease inhibitor cocktail (Roche; Cat# 4693132001). Protein concentration was determined via Bradford assay (Bio-Rad). Protein solution was adjusted to equal concentration and incubated for 20 min at 40°C in SDS-sample buffer (2% SDS, 10% glycerol, 75 mM Tris-HCl pH 6.8, 0.02% bromphenol blue, 5% beta-mercaptoethanol). SDS-PAGE gels were prepared with acrylamide/bisacrylamide (ratio 97%/3%) concentrations of 4% for the stacking and 10% for the separating gel. Protein size was determined via Sea-Blue Plus 2 pre-stained protein standards (Invitrogen; Cat# LC5925). Immunoblotting of PAGEs was conducted onto Amersham nitrocellulose or PVDF membranes (GE healthcare) for 2 h at 1.6 mA/cm^2^ using blotting buffer (25 mM Tris, 185 mM glycine, 20% methanol). Membranes were blocked in 5% milk powder in TBS + 0.05% Tween-20.

### Cell treatments

Protein stability was followed by treatment with cycloheximide (Sigma; Cat# C4859) at a concentration of 100 µm*ml^-1^ over the course of 8 h. At indicated time points, cells were collected and lysed according to the protocol above. Cell viability was assessed after treatment with actinomycin D (Sigma, Cat# A9415; 1µM), venetoclax (Selleckchem; Cat# S8048; 1µM), A-1155463 (Selleckchem; Cat# S7800; 1µM), staurosporine (Sigma, Cat# S6942; 1µM) or thapsigargin (Sigma; Cat# T9033; 2µM) for 16 h. ATP-synthase was inhibited by oligomycin treatment (Sigma, Cat# O4876; 10µM) for 18 h.

### Mitochondrial isolation

Mitochondrial isolation was conducted from cell pellets using isolation buffer (220 mM mannitol, 70 mM sucrose, 2 mM EGTA, 20 mM HEPES-KOH pH 7.4, 1 mM PMSF (Roche, Cat# 10837091001)), complete protease inhibitor cocktail (Roche, Cat# 52434800), 0.1% BSA) including a 15 min swelling step. Cells were disrupted using 15 strokes in a glass mortar and PTFE pestle at 1000 rpm with a rotating homogenizer (Schuett biotec, Cat# 3201011). The mitochondrial containing fraction was separated at 1000 × g for 15 min and subsequently pelleted at 8000 × g for 15 min before washing in isolation buffer without BSA. Freshly isolated mitochondria were used for experiments relying on an intact mitochondrial membrane potential.

### Immunoprecipitation

Isolated mitochondria were incubated for 15 min in lysis buffer (20 mM HEPES-NaOH pH7.5, 300 mM NaCl) containing 4 g digitonin per g mitochondria. Centrifuged lysate was incubated with washed EZview Red anti-FLAG M2 affinity gel (Millipore, Cat# F2426) for 1 h. Beads were washed (10 mM HEPES-NaOH pH7.5, 150 mM NaCl, 0.1% Triton X-100) three times before analyzing via SDS-PAGE or mass spectrometry-based proteomics.

### Fluorescence-activated cell sorting (FACS)

#### NucView 488 RedDot 2

HeLa cells were plated 20*10^4^ each 6 well. 24 h later they were treated with the staurosporine (0.1 μM) for 16 h. Cells were detached using trypsin, resuspended at a density of 10^6^ cells/ml in culture medium that contain NucView 488 substrate (2.5 μM) and RedDot 2 (0.25X). Cells were incubated at room temperature for 30 minutes protected from light, resuspended in PBS and analyze by flow cytometry. NucView fluorescence was measured in the green detection channel (ex488/em530 nm). RedDot 2 fluorescence in the far-red channel (ex640/em780 nm).

### MitoSOX

HeLa cells were plated 20*10^4^ each 6 well. 24 h later were detached using trypsin, resuspended at a density of 10^6 cells/ml in PBS containing MitoSOX Red for 30 min and then analyzed using flow cytometry. Rotenone (0.5 µM) was added during MitoSox Red staining.

### EM

Cells were fixed in fixation buffer (2% glutaraldehyde, 2.5% sucrose, 3 mM CaCl2, 100 mM HEPES-KOH pH 7.2) at RT for 30 min and 4°C for 30 min. Samples were washed three times in 1% osmium tetroxide, 1.25% sucrose and 1% potassium ferrocyanide in 0.1 M sodium cacodylate buffer. After dehydration in alcohol gradient series and propylene oxide, the tissue samples were embedded. Ultrathin sections were cut on a diamond knife (Diatome, Biel, Switzerland) on a Leica ultramicrotome, place on copper grids (Science Services, 100mesh). Sections were stained with uranyl acetate (Plano, 1.5%) and lead citrate (Sigma) and examined with an electron microscope (JEM 2100 Plus, JEOL) with an OneView 4K camera (Gatan) with DigitalMicrograph software at at 80 kV.

### Calcium retention capacity

Isolated mitochondria from HEK293T cells were resuspended in assay buffer (100 mM sucrose, 4 mM MgCl2, 80 mM KCl, 5 mM Na-succinate, 20 mM HEPES-KOH pH7.6, 0.4 mM ADP, 0.5 mM NADH, 0.01 mM EGTA, 0.5 µM Calcium Green 5N (Thermofisher), 2 mM KH2PO4) to a concentration of 1 µg*µl^-1^. Mitochondria were subjected to 10 µl calcium chloride injections until disruption. Fluorescence was measured in a Perkin Elmer EnVision 2105 plate reader at excitation 485 nm and emission 53 5nm.

### Protein expression and purification

Codon-optimized DNA sequence for expression in *E. coli* of the human TMBIM5 protein was synthesized from PCR fragments by GeneArt (Life Technologies) and subsequently cloned into the bacterial overexpression vector pMALp2x (NEB). The recombinant TMBIM5 protein expression was performed in BL21(DE3) *E. coli*. The bacteria were propagated in TB medium. Protein expression was induced with 0.5 mM IPTG and was carried out at 37°C for 4 h in an orbital shaker. Bacterial cells were harvested at 3000 × g for 20 min, washed in 200 mM NaCl, 5% glycerol, 1 mM EDTA, 20 mM HEPES-KOH pH7.4 and stored at -20°C. Bacterial pellets were resuspended in buffer (200 mM NaCl, 5% glycerol, 1 mM EDTA, 1 mM DTT, 20 mM HEPES-KOHH pH7.4) which was supplemented with protease inhibitor cocktail (Roche) and lysed in an Emulsiflex. The obtained cell lysate was supplemented with 0.05% DDM and spun down at 180 000 × g at 4°C for 20 min. The supernatant was applied onto MBPTrap affinity column (Cytiva; Cat# 28918780) and the bound fraction was subsequently eluted with 10 mM maltose. Protein-containing fractions were subjected to Factor Xa digest (NEB). Resulting mixture of untagged TMBIM5 and free MBP-tag was separated on HiLoad Superdex 16/600 75 pg column equilibrated in 120 mM NMDG, 1mM DTT, 0.05% DDM, 10 m MOPS-KOH pH 7.4.

### Preparation of TMBIM5-proteoliposomes

All lipids used for preparation of liposomes were purchased from Avanti Polar Lipids. The lipid mixture mimicking the composition of the mitochondrial inner membrane contained 45% L-*α*-phosphatidylcholine, 20% L-*α*-phosphatidylethanolamine, 15% L-*α*-phosphatidyl- inositol, 15% cardiolipin and 5% L-*α*-phosphatidylserine. Lipid stocks prepared in methanol/ chloroform mixture were dried under constant nitrogen flow. Dried lipid films were re- hydrated in 120 mM NMDG, 10 mM MOPS pH 7.4 buffer supplemented with 100 μM of Fura- 2 (Invitrogen, Cat# F1200) for 10 min and subjected to the sequence of freeze-and-thaw cycles of ten repetitions. The obtained lipid vesicles were extruded through a PVDF filter with pore diameter of 400 nm and used for protein incorporation. Prior to protein incorporation, Fura-2- containing liposomes were partially solubilized with 0.1% DDM and incubated for 30 min at RT under gentle shacking. Purified TMBIM5 protein was added to the pre-solubilized liposomes in a 100 to 1 (w/w) ratio and incubated for 1 h to allow spontaneous protein insertion. The incorporation process was promoted by removing the detergent from the mixture using adsorbent BioBeads SM-2 (BioRad). Formed proteoliposomes were separated from the BioBeads and the proteoliposomes were harvested by centrifugation at 180 000 × g for 20 min at 4°C, washed with 120 mM NMDG, 10 mM MOPS-KOH pH8 and used for Ca^2+^-flux measurements.

Ca^2+^ flux was initiated by adding CaCl2 (100 μM). The fluorescence intensity was monitored every 1 s in a Perkin Elmer EnVision 2105 plate reader at excitation 340 nm and 380 nm, emission 510 nm. Ionomycin (10 mM) was added at the end of the experiment for calibration. GdCl3 (100 μM) was added immediately before the experiment were indicated.

### Electrophysiological characterization of TMBIM5

Electrophysiological characterization of TMBIM5 was carried out using the planar lipid bilayer technique (Denkert et al., 2017; Vasic et al., 2020). Electrical current recordings were performed using Ag/AgCl electrodes in glass tubes, embedded in a 2 M KCl agar-bridge. The electrode in the trans chamber was connected to the headstage (CV-5-1GU) of a Geneclamp 500B current amplifier (Molecular Devices, CA, USA) and acted as a reference electrode. Data were acquired using a Digidata 1440A A/D converter and recorded using the AxoScope 10.3 and Clampex 10.3 software (Molecular Devices). Data analysis was carried out using OriginPro 8.5G (OriginLab, MA, USA).

TMBIM5-proteoliposomes were inserted into the planar lipid bilayer by osmotic driven fusion in asymmetric buffer condition with 250 mM KCl, 10 mM MOPS, pH 7.0 in the cis chamber and 20 mM KCl, 10 mM MOPS, pH 7.0 in trans chamber. After incorporation of TMBIM5 into the planar lipid bilayer, both of the chambers were perfused with 250 mM CaCl2, 10 mM MOPS, pH 7.0 to attain symmetrical buffer conditions and voltage-clamp recordings as well as voltage-ramps were performed. To investigate the inhibitory effects of Gd^3+^ ions on TMBIM5, 9 mM GdCl3 was added to both cis and trans chambers and stirred for 2 min before the current recordings. Voltage ramps were recorded from -80 mV to +80 mV and the current voltage relationship was plotted. Current traces were recorded at constant voltage for one minute. The open probability (Popen) was calculated from the maximum current recorded, divided by the mean current using the data from three experimental replicates.

### Proteomics

#### Determination of the AFG3L2 interactome by data dependent acquisition

This method part is related to Figure 1A and Suppl. Table 01.

#### Protein Digestion

Proteins bound to the EZview Red anti-FLAG M2 affinity gel (Millipore; catalog number F2426) beads were lysed using 60 µL of 4% SDS in 100 mM HEPES pH 8.5 at 70°C for 10 min on a ThermoMixer with shaking (550 rpm). Proteins were reduced (10 mM TCEP) and alkylated (20 mM CAA) in the dark for 45 min at 45°C and processed via the SP3 digestion protocol (Hughes et al., 2019). Washed SP3 beads (SP3 beads (Sera-Mag(TM) Magnetic Carboxylate Modified Particles (Hydrophobic, catalog number GE44152105050250), Sera-Mag(TM) Magnetic Carboxylate Modified Particles (Hydrophilic, catalog number GE24152105050250) from Sigma Aldrich) were mixed equally, and 3 µL of bead slurry were added to each sample. Acetonitrile was added to a final concentration of 50% and washed twice using 70% ethanol (V=200 µL) on an in-house made magnet. After an additional acetonitrile wash (V=200 µL), 5 µL 10 mM HEPES pH 8.5 containing 0.5 µg trypsin (Sigma, #T6567- 1MG) and 0.5 µg LysC (Wako, # 129-02541) were added to each sample and incubated overnight at 37°C. Peptides were desalted on a magnet using 2 x 200 µL acetonitrile. Peptides were eluted in 10 µL 5% DMSO in LC-MS water (Sigma Aldrich) in an ultrasonic bath for 10 min. Samples were dried and stored at -20°C until subjected to LC-MS/MS analysis. Peptides were then reconstituted in 10 µL 2.5% formic acid and 2% acetonitrile and 3 µL were used for a LC-MS/MS run.

#### Liquid Chromatography and Mass Spectrometry

LC-MS/MS instrumentation consisted out of an Easy-LC 1200 (Thermo Fisher Scientific) coupled via a nano-electrospray ionization source to an QExactive HF-x mass spectrometer (Thermo Fisher Scientific). For peptide separation an in-house packed column (inner diameter: 75 µm, length: 40 cm) was used. A binary buffer system (A: 0.1 % formic acid and B: 0.1 % formic acid in 80% acetonitrile) was applied as follows: Linear increase of buffer B from 4% to 32% within 33 min, followed by a linear increase to 55% within 5 min. The buffer B content was further ramped to 95% within 2 min. 95 % buffer B was kept for further 5 min to wash the column. Prior each sample, the column was washed using 6 µL buffer A and the sample was loaded using 7 µL buffer A.

The mass spectrometer operated in a data-dependent mode and acquired MS1 spectra at a resolution of 60000 (at 200 m/z) using a maximum injection time of 20 ms and an AGC target of 3e6. The scan range was defined from 350-1650 m/z and data type was set to profile. MS2 spectra were acquire at a 15000 resolution (at 200 m/z) using an isolation window of 1.4 m/z and a normalized collision energy of 32. The Top22 peaks were targeted for MS2 spectra acquisition. The first mass was set to 110 m/z. Dynamic exclusion was enable and set to 20 s.

#### Data Analysis of Data Dependent Acquisition Proteomics

The Andromeda Search Engine implemented into the MaxQuant software (1.6.0.1) was used to analyzed DDA files to investigate the AFG3L2 interactome. MS/MS spectra were correlated against the UniProt reference proteome (downloaded Nov. 2019, containing 20362 protein amino acid sequences) using default settings. The match-between runs algorithm was enabled as well as the label free quantification feature (min. count = 1). The protein groups output file was then further processed in Perseus by log2 transforming the LFQ intensities. Protein groups were then filtered for being quantified in 5 out of 5 replicates followed by imputation of missing values from a down-shifted (downshift: 1.8*standard dev., width: 0.4* standard dev.) gaussian distribution. To identify potential interactors, we used a two-sided t-test following by permutation-based FDR estimation to correct for multiple testing (FDR < 0.05). Gene Ontology annotations were added and data were filtered solely for visualization based on the localization to the mitochondria using the GO cellular component information.

#### Whole cell proteomics using data independent acquisition

This method part is related to Figure 4G-I, Figure 5A-C, Figure 6A-B, Figure 7A-B, Suppl Table 2.

#### Protein Digestion

40 µL of 4% SDS in 100 mM HEPES pH 8.5 was pre-heated to 70°C and added to the cell pellet for further 10 min incubation at 70°C on a ThermoMixer (shaking: 550 rpm). The protein concentration was determined using the 660 nm Protein Assay (ThermoFisher Scientific, catalog number 22660). 50 µg of protein was subjected for tryptic digestion. Proteins were reduced (10 mM TCEP) and alkylated (20 mM CAA) in the dark for 45 min at 45 °C. Samples were subjected to SP3 based digestion as described above (Hughes *et al*., 2019). Generated peptides were eluted from magnetic beads in 10 µL 5% DMSO in LC-MS water (Sigma Aldrich) in an ultrasonic bath for 10 min. Samples were then dried and stored at -20°C until subjected to LC-MS/MS analysis. Peptides were then reconstituted in 10 µL 2.5% formic acid and 2% acetonitrile and 3 µL were used for a LC-MS/MS run.

#### Liquid Chromatography and Mass Spectrometry

LC-MS/MS instrumentation consisted out of an Easy-LC 1200 (Thermo Fisher Scientific) coupled via a nano-electrospray ionization source to an Exploris 480 or QExactive HF-x mass spectrometer (Thermo Fisher Scientific). For peptide separation an in-house packed column (inner diameter: 75 µm, length: 40 cm) was used. A binary buffer system (A: 0.1 % formic acid (Thermo Fisher Scientific, cat# LS118-1) and B: 0.1 % formic acid in 80% acetonitrile (Thermo Fisher Scientific, cat# LS122)) was applied as follows: Linear increase of buffer B from 4% to 27% within 69 min, followed by a linear increase to 45% within 5 min. The buffer B content was further ramped to 65% within 5 min and then to 95% within 6 min. 95% buffer B was kept for further 10 min to wash the column. Prior each sample, the column was washed using 5 µL buffer A and the sample was loaded using 8 µL buffer A.

The RF Lens amplitude was set to 55%, the capillary temperature was 275°C and the polarity was set to positive. MS1 profile spectra were acquired using a resolution of 60,000 (at 200 m/z) at a mass range 320-1150 m/z and an AGC target of 1 × 106.

For data independent MS/MS spectra acquisition, 48 windows were acquired at an isolation m/z range of 15 Th and the isolation windows overlapped by 1 Th. The fixed first mass was to 200. The isolation center range covered a mass range of 350–1065 m/z. Fragmentation spectra were acquired at a resolution of 15,000 at 200 m/z using a maximal injection time of 22 ms and stepped normalized collision energies (NCE) of 26, 28, 30. The default charge state was set to 3. The AGC target was set to 900% (Exploris 480) or 1e6 (QExactive HF-x). MS2 spectra were acquired as centroid spectra.

### Data Analysis

For the analysis of DIA (Data independent acquisition) experiments, expect pulseSILAC (see below), we utilized DIA-NN version 1.7.15 (Demichev et al., 2020). The library-free approach was used based on the human Uniport reference proteome (downloaded Nov. 2019, containing 20362 protein amino acid sequences) which predicts MS2 spectra using neuronal network. The deep-learning option was enabled. Quantification strategy was set to ‘robust LC (high accuracy)’. The precursor range was adjusted to 330 – 1200 m/z matching the acquired mass range. The RT profiling option was enabled. Otherwise, default settings were used. To identify significantly different proteins, a two-sided t-test was applied using log2 transformed MaxLFQ intensities calculated by DIA-NN (Cox et al., 2014). The FDR was controlled to 5% using a permutation-based approach in the Perseus software (Tyanova et al., 2016). If applicable, a two/three ANOVA was calculated in the Perseus software. Raw output of DIA-NN analysis as well as raw files were deposited to the PRIDE repository.

#### CHX-chase experiment

Log2 transformed LFQ intensities were utilized to determine the degradation rate of each protein by fitting a first-order kinetic. The area under curve was determined using trapezoid rule in the software InstantClue (Nolte et al., 2018). The AUC values were log transformed and a two-sided t-test was utilized to identify significantly regulated degradation rates.

#### pulse SILAC data

To analyze pulseSILAC DIA data, a library was first created using acquired DIA files Spectronaut’s Pulsar search engine using exclusively y-ions but otherwise default settings. Then in a second analysis, the individual files were analyzed using the generated spectra library. Data were exported on the precursor level and were transformed as follows. First, the remaining heavy fraction was determined by calculating the natural logarithm of H/L / (H/L + 1). Second, the data were adjusted by calculation the median of the remaining fraction for each group (e.g. timepoint and genotype) and individual replicates were adjusted to the overall group median. Then, data were filtered by removing values that were found to be outliers (1.8 x inter quantile range) which were replaced by the group mean. A linear model was fit to the data and the r-value, p-value, slope and intercepts as well as their standard errors were calculated for each individual replicate (time point 48 h was removed since it was found to be outside the linear range). Slope values with a r-value above -0.75 were removed by replacement with NaN. To compare the slopes between genotypes, a two-sided t-test was performed. Analysis was performed using the Instant Clue (v 0.11.1) software suite (www.instantclue.de) (Nolte *et al*., 2018).

### Quantification and statistical analysis

Densitometry data for western blots were generated using Fiji. Representative images of at least three independent experiments are shown. Graphs were plotted using Prism (GraphPad). Volcano plots and heatmaps were generated with InstantClue. Error bars represent standard deviation of the mean. Statistical significance of the data was mostly assessed by using Student’s t test, comparing control and test conditions, and it is described in the figure legends.

