## Supplemental Figures for "Regulation of mitochondrial proteostasis by the proton gradient"

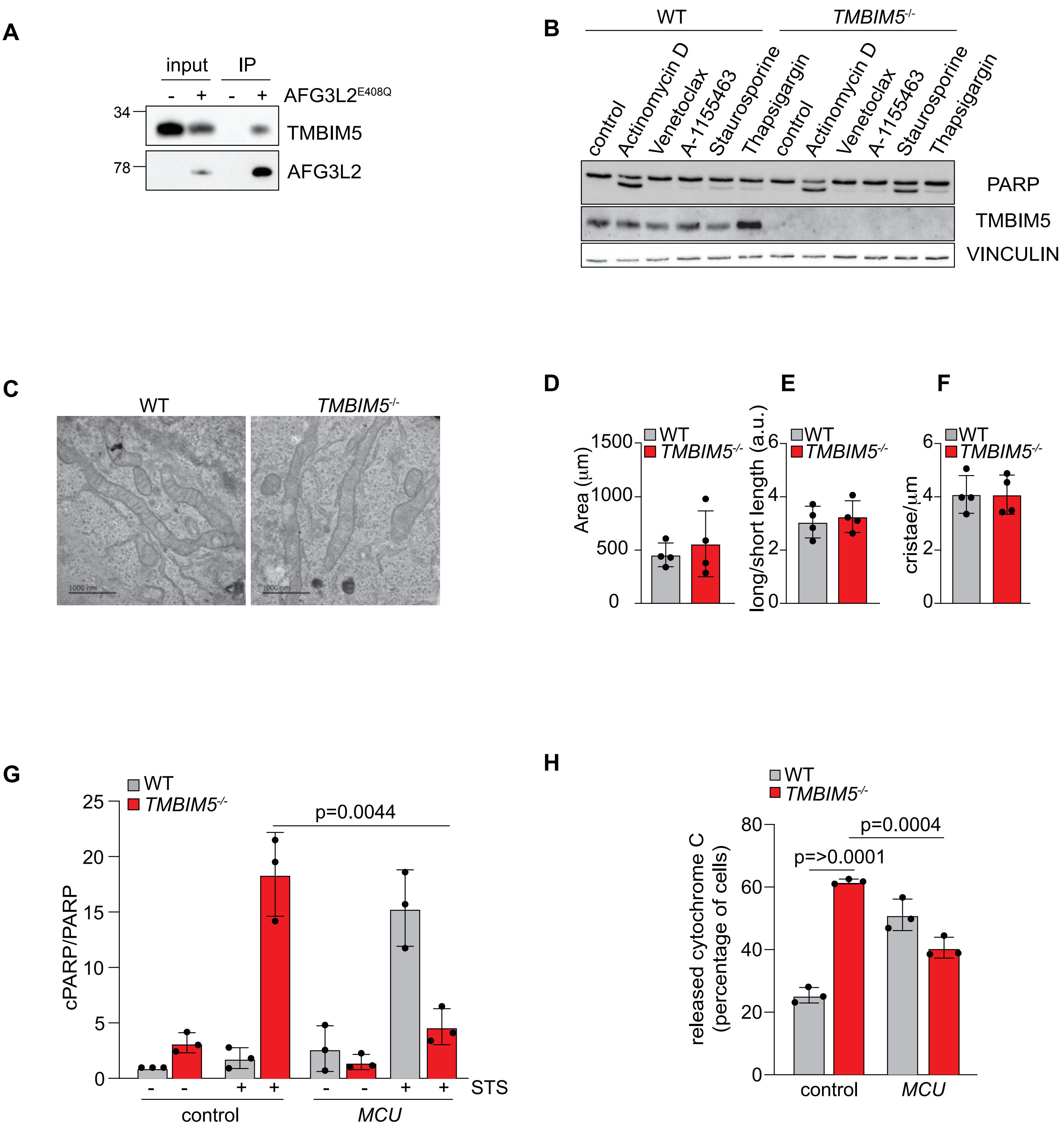
A
input
IP
AFG3L2E408Q
34
78
TMBIM5
AFG3L2
B
WT
TMBIM5-/-
control
Actinomycin D
Venetoclax
A-1155463
Staurosporine
Thapsigargin
control
Actinomycin D
Venetoclax
A-1155463
Staurosporine
Thapsigargin
PARP
TMBIM5
VINCULIN
C
WT
TMBIM5-/-
1000 nm
1000 nm
D
E
F
WT
TMBIM5-/-
Area (µm)
1500
1000
500
0
long/short length (a.u.)
cristae/µm
6
4
2
0
G
WT
TMBIM5-/-
cPARP/PARP
25
20
15
10
5
0
p=0.0044
STS
control
MCU
H
WT
TMBIM5-/-
released cytochrome C (percentage of cells)
80
60
40
20
0
p=>0.0001
p=0.0004
control
MCU

**Fig. S1**

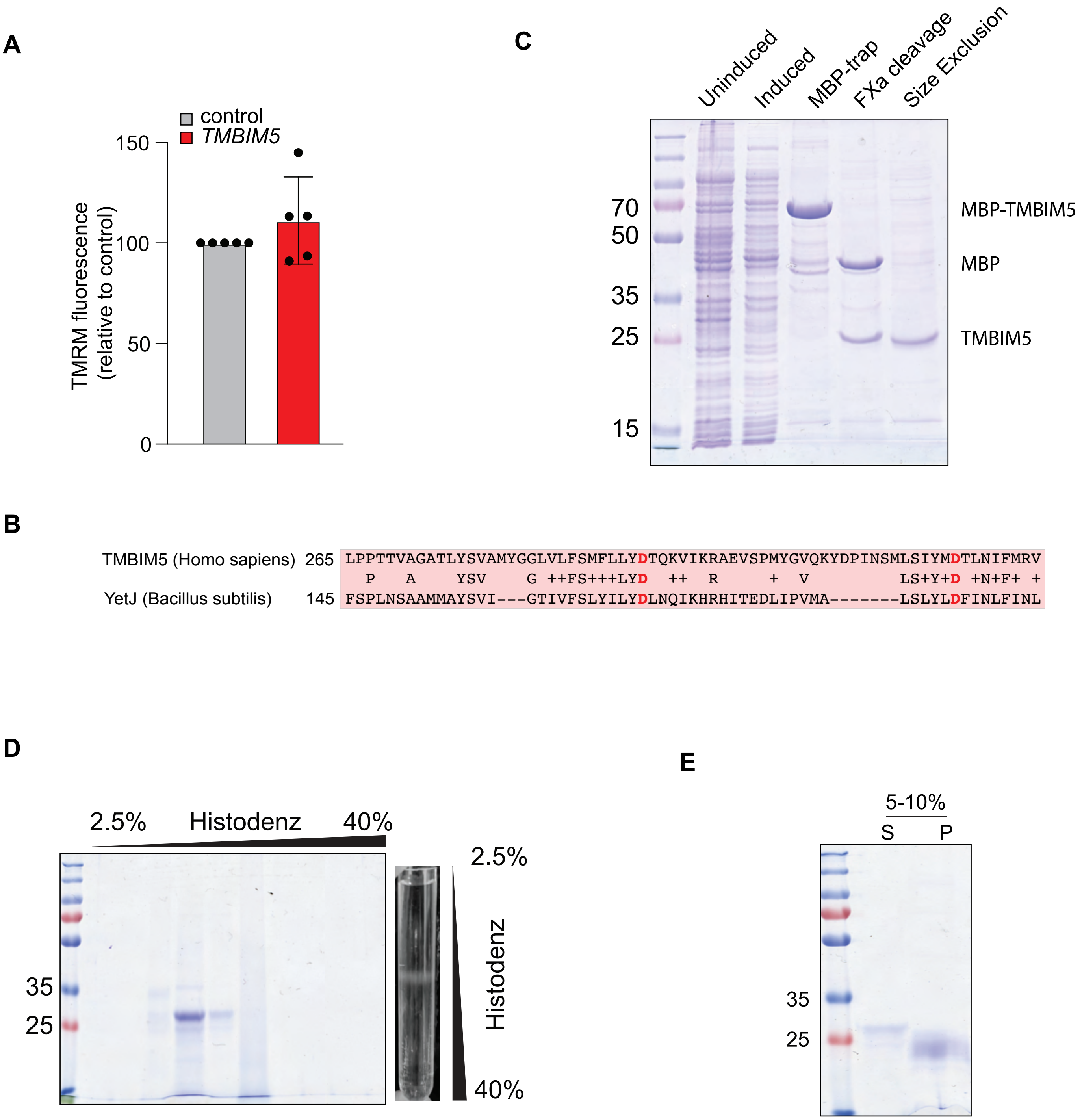
A
control
TMBIM5
TMRM fluorescence
(relative to control)
150
100
50
0
C
Uninduced
Induced
MBP-trap
FXa cleavage
Size Exclusion
70
50
35
25
15
MBP-TMBIM5
MBP
TMBIM5
B
TMBIM5 (Homo sapiens) 265 LPPTTVAGATLYSVAMYGGLVLFSMFLLYDTQKVIKRAEVSPMYGVQKYDPINSMLSIYMDTLNIFMRV
P A YSV G ++FS+++LYD ++ R + V LS+Y+D +N+F+ +
YetJ (Bacillus subtilis) 145 FSPLNSAAMMAYSVI---GTIVFSLYILYDLNQIKHRHITEDLIPVMA-------LSLYLDFINLFINL
D
2.5%
Histodenz
40%
35
25
2.5%
Histodenz
40%
E
5-10%
S
P
35
25

**Fig. S2**

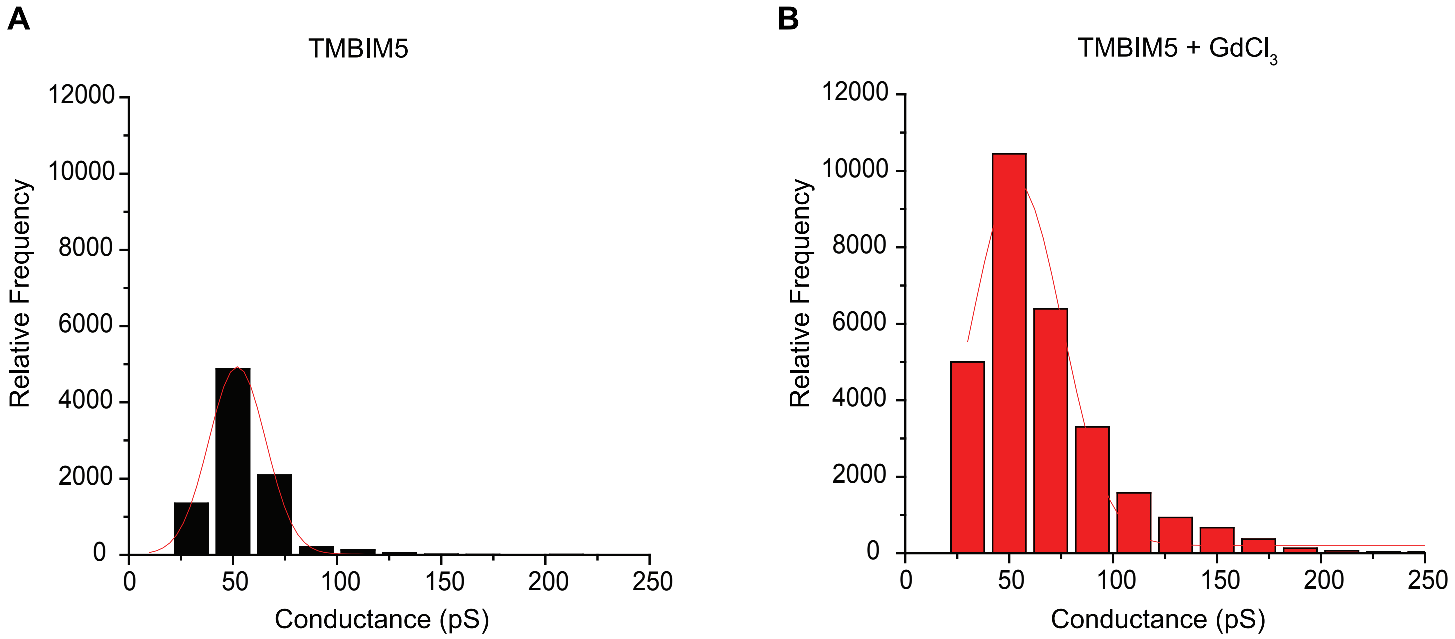
A
TMBIM5
12000
10000
8000
6000
4000
2000
0
Relative Frequency
0
50
100
150
200
250
Conductance (pS)
B
TMBIM5 + GdCl3
12000
10000
8000
6000
4000
2000
0
Relative Frequency
0
50
100
150
200
250
Conductance (pS)

**Fig. S3**

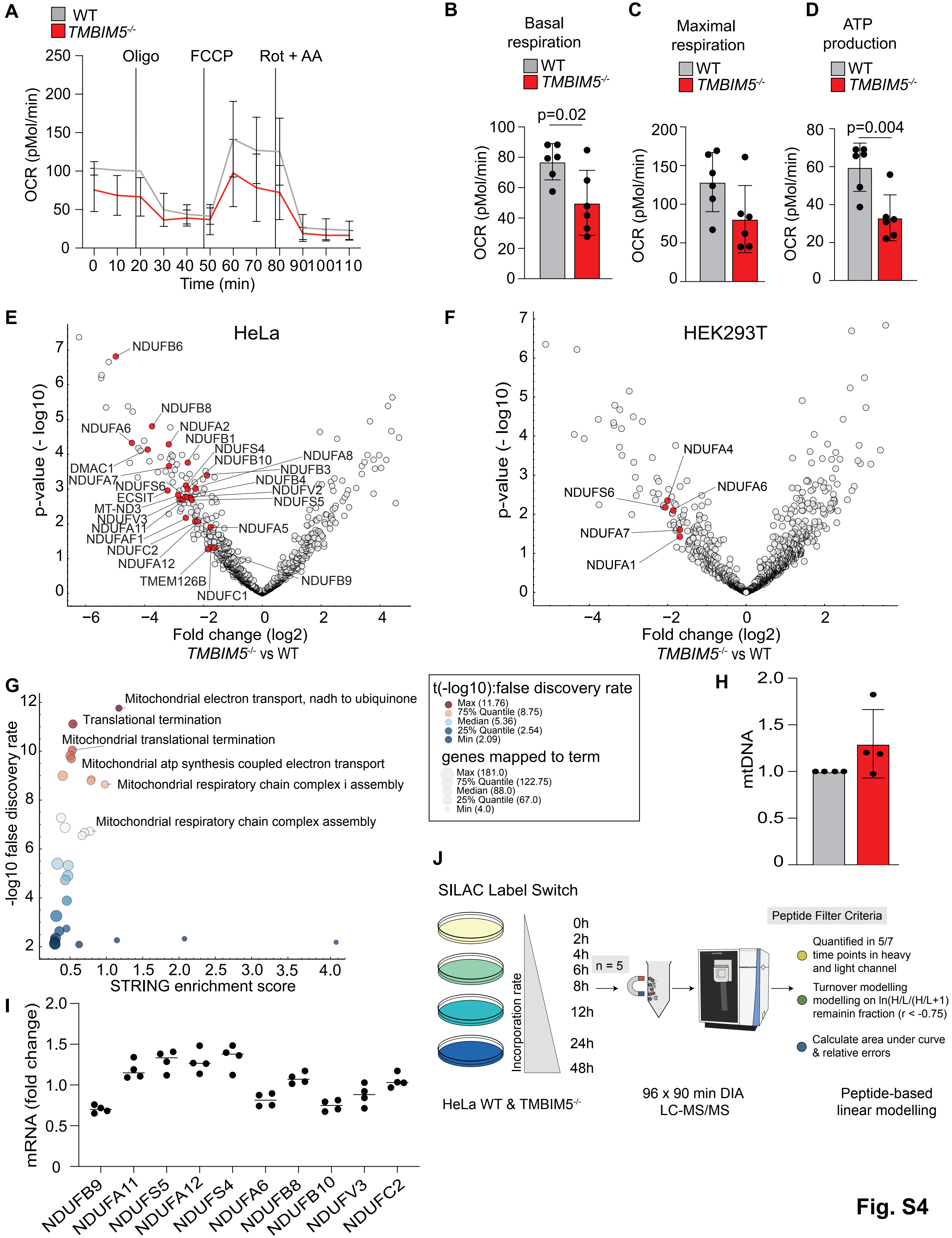

A
WT
TMBIM5-/-
Oligo
FCCP
Rot + AA
OCR (pMol/min)
Time (min)
B
Basal respiration
p=0.02
C
Maximal respiration
D
ATP production
p=0.004
E
HeLa
p-value (- log10)
Fold change (log2)
TMBIM5-/- vs WT
NDUFB6
NDUFB8
NDUFA6
NDUFA2
NDUFB1
NDUFS4
NDUFB10
NDUFA8
NDUFB3
DMAC1
NDUFA7
NDUFS6
NDUFB4
NDUFV2
ECSIT
NDUFS5
MT-ND3
NDUFV3
NDUFA11
NDUFAF1
NDUFC2
NDUFA12
TMEM126B
NDUFC1
NDUFA5
NDUFB9
F
HEK293T
NDUFA4
NDUFS6
NDUFA6
NDUFA7
NDUFA1
G
-log10 false discovery rate
STRING enrichment score
Mitochondrial electron transport, nadh to ubiquinone
Translational termination
Mitochondrial translational termination
Mitochondrial atp synthesis coupled electron transport
Mitochondrial respiratory chain complex i assembly
Mitochondrial respiratory chain complex assembly
t(-log10):false discovery rate
Max (11.76)
75% Quantile (8.75)
Median (5.36)
25% Quantile (2.54)
Min (2.09)
genes mapped to term
Max (181.0)
75% Quantile (122.75)
Median (88.0)
25% Quantile (67.0)
Min (4.0)
H
mtDNA
I
mRNA (fold change)
NDUFB9
NDUFA11
NDUFS5
NDUFA12
NDUFS4
NDUFA6
NDUFB8
NDUFB10
NDUFV3
NDUFC2
J
SILAC Label Switch
0h
2h
4h
6h
8h
12h
24h
48h
n = 5
Incorporation rate
Peptide Filter Criteria
Quantified in 5/7 time points in heavy and light channel
Turnover modelling modelling on ln(H/L/(H/L+1)) remainin fraction (r < -0.75)
Calculate area under curve & relative errors
HeLa WT & TMBIM5-/-
96 x 90 min DIA LC-MS/MS
Peptide-based linear modelling


**Fig. S4**

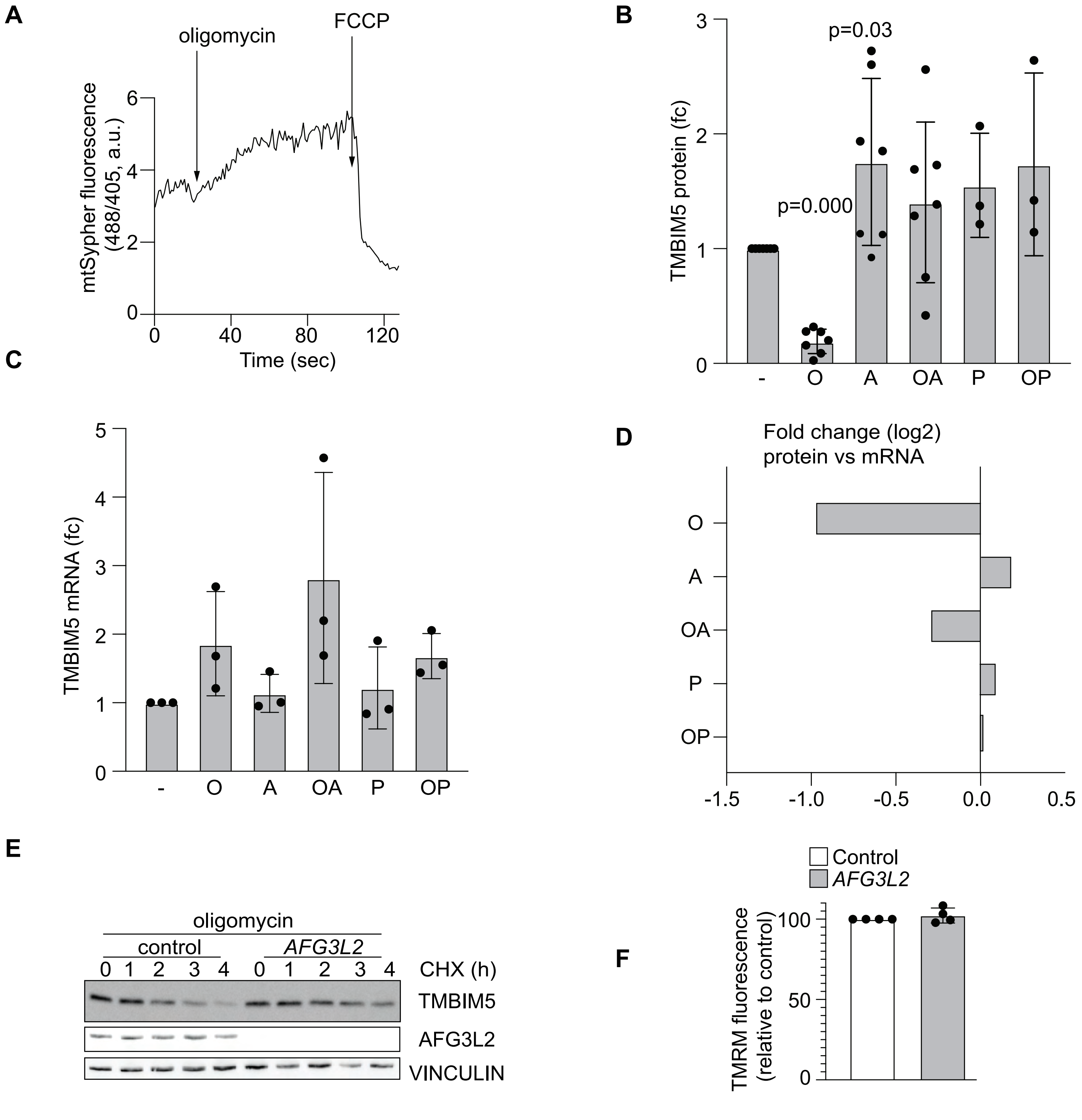
A
oligomycin
FCCP
mtSypher fluorescence (488/405, a.u.)
Time (sec)
B
TMBIM5 protein (fc)
p=0.03
p=0.000
- O A OA P OP
C
TMBIM5 mRNA (fc)
- O A OA P OP
D
Fold change (log2)
protein vs mRNA
O
A
OA
P
OP
E
oligomycin
control
AFG3L2
CHX (h)
TMBIM5
AFG3L2
VINCULIN
F
Control
AFG3L2
TMRM fluorescence (relative to control)

Fig. S5

**Supplementary Figures**

**Figure S1. Mitochondrial $Ca^{2+}$ overload triggers apoptotic death of *TMBIM5*$^{-/-}$ cells**

**Related to Figure 1**

**(A)** Representative immunoblot of immunoprecipitates using anti-FLAG M2 beads of mitochondrial lysates which were isolated from Flp-In HEK293T-REx cells expressing AFG3L2$^{E408Q\ FLAG}$ when indicated.

**(B)** Representative immunoblot of WT and *TMBIM5*$^{-/-}$ HeLa cells treated with the indicated drugs for 16 h. Control (0.1% DMSO), actinomycin D (1 µM), venetoclax (1 µM), A-1155463 (1 µM), staurosporine (1 µM), thapsigargin (2 µM). (n=3 independent experiments).

**(C)** Representative transmission electron microscopy images of WT and *TMBIM5*$^{-/-}$ HeLa cells. Mitochondrial morphology is shown. Scale bar 1000 nm. (n=4 independent experiments).

**(D)** Quantification of the electron microscopy images of HeLa WT and *TMBIM5*$^{-/-}$ cells. Mitochondrial parameters (area, length and cristae) were calculated with FiJi.

**(E)** Quantification of the electron microscopy images of WT and *TMBIM5*$^{-/-}$ HeLa cells as in Figure S1F. The ratio between the long length and the short length of mitochondria is shown.

**(F)** Quantification of electron microscopy images of WT and *TMBIM5*$^{-/-}$ HeLa cells as in Figure S1F. Number of cristae per mitochondrial length (µM) is shown.

**(G)** Quantification of Figure 1C. The ratio between cleaved PARP (cPARP; 89 kDa) and PARP (116 kDa). WT and *TMBIM5*$^{-/-}$ HeLa cells transfected with scrambled siRNA (control) or siRNA targeting *MCU* for 72 h. When indicated, samples were treated with staurosporine (STS; 1 µM) for 16 h. (n=3 independent experiments).

**(H)** Quantification of WT and *TMBIM5*$^{-/-}$ HeLa cells with cytoplasmic cytochrome c. After transfection with scrambled siRNA (control) or siRNA targeting *MCU* for 72 h cells were incubated for 16 h with staurosporine (0.1 µM) in the presence of Z-VAD-FMK (50 µM) and epoxomicin (1 µM) to prevent apoptosis. Cells were analyzed by immunofluorescence microscopy using antibodies directed against cytochrome c (green) and TOMM20 (magenta). (n=3 independent experiments; number of cells > 150 in each experiment).

**Figure S2. Reconstitution of TMBIM5 into liposomes**

**Related to Figure 2**

**(A)** Mitochondrial membrane potential was monitored by TMRM staining in WT HeLa cells transfected with scrambled siRNA (control) or siRNA targeting *TMBIM5* for 72 h. Fluorescent intensity was calculated relative to the value upon CCCP addition (15 µM). Values are expressed as mean relative to control. Measurements of 5 different preparations were performed in triplicate.

**(B)** Sequence alignment of TMBIM5 (Homo sapiens) and YetJ (*Bacillus subtilis*). Conserved amino acids between the two species are written in the middle line. We used red color to highlight the amino acids forming the di-aspartyl dyad.

**(C)** Representative Coomassie-stained SDS-polyacrylamide gel (12.5%) demonstrating enrichment of recombinant TMBIM5 protein upon purification.

**(D)** Flotation assay demonstrating co-migration of recombinant TMBIM5 protein with the liposomes after ultracentrifugation in a discontinuous histodenz gradient. A representative Coomassie-stained SDS-polyacrylamide gel (12.5%) is shown. The ultracentrifugation tube containing TMBIM5-liposomes in discontinuous histodenz gradient after ultracentrifugation but before fractionation is shown on the right.

**(E)** Sodium bicarbonate extraction. Supernatant (S) and pellet (P) fractions of TMBIM5-proteoliposomes after treatment with 0.1 M sodium bicarbonate (pH 11.5) analyzed by SDS-polyacrylamide gel (12.5%) electrophoresis. Pellet (P), and supernatant (S) are indicated in the figure. The presence of TMBIM5 in the pellet fraction indicates its integration into the lipid bilayer.

**Figure S3. TMBIM5 acts as a $Ca^{2+}/H^{+}$ exchanger.**

**Related to Figure 3**

**(A)** TMBIM5 conductance state histogram in $CaCl_2$ buffer. (n=3 independent experiments).

**(B)** TMBIM5 conductance state histogram in $CaCl_2$ buffer containing $GdCl_2$. (n=3 independent experiments).

**Figure S4. TMBIM5 controls oxidative phosphorylation**

**Related to Figure 4**

**(A)** Oxygen consumption rate (OCR) of WT and *TMBIM5*$^{-/-}$ HEK293T cells in glucose media. Traces are mean ± SD of 6 independent experiments, each one run at least in triplicate. Labeled lines denotes injections of oligomycin (Oligo, 2 μM), FCCP (0.5 μM), rotenone and antimycin A (Rot + AA; both 0.5 μM).

**(B)** Basal respiration calculated from OCR experiment of WT and *TMBIM5*$^{-/-}$ HEK293T cells in (A). Two-tailed *t*-test. A p-value of <0.05 was considered statistically significant.

**(C)** Maximal respiration calculated from OCR experiment of WT and *TMBIM5*$^{-/-}$ HEK293T cells in (A).

**(D)** ATP production calculated from OCR experiments of WT and *TMBIM5*$^{-/-}$ HEK293T cells in (A). Two-tailed *t*-test. A p-value of <0.05 was considered statistically significant.

**(E)** Volcano plots of mitochondrial protein changes in *TMBIM5*$^{-/-}$ HeLa cells when compared to WT HeLa cells. Significantly enriched mitochondrial complex I proteins at an FDR cutoff of 0.05 are colored in red. (n=6 independent experiments). P-values were calculated from a two-sided t-test followed by a permutation-based FDR controlling to 0.05. See also Table S2.

**(F)** Volcano plots of mitochondrial protein changes in *TMBIM5*$^{-/-}$ HEK293T cells when compared to WT HEK293T cells. Significantly enriched mitochondrial complex I proteins at an FDR cutoff of 0.05 are colored in red. (n=5 independent experiments). P-values were calculated from a two-sided t-test followed by a permutation-based FDR controlling to 0.05. See also Table S3.

**(G)** Scatter plot comparing the STRING enrichment score versus the -log10 transformed false discovery rate (color encoded). The number of genes mapped to the enriched GOBP term is encoded by size.

**(H)** mtDNA levels from HeLa cells assessed by qPCR amplification of mitochondrial CYTB (n=4 independent experiments).

**(I)** Expression of complex I subunits monitored by RT-qPCR. (n=4 independent experiments).

**(J)** Workflow of SILAC chase experiments in WT and *TMBIM5*$^{-/-}$ HeLa cells.

**Figure S5.**

**Related to Figure 6**

**(A)** Representative Sypher3mito traces of HeLa cells treated with oligomycin (10 μM) or FCCP (10 μM) at the indicated timepoints. Ratios of the absorbance at 488 nm and 405 nm are shown in arbitrary units (a.u.).

(B) Quantification of Figure 6C. WT HeLa cells treated with the indicated drugs for 16 h. antimycin A (A; 10 μM), piericidin (P; 10 μM) and oligomycin (O; 10 μM). (n=7 independent experiments).

**(C)** Quantification of *TMBIM5* mRNA monitored by RT-qPCR of HeLa cells treated as in Figure 6C. (n=3 independent experiments).

(D) Bar graph displaying log2 fold changes between transcript (mRNA) (Figure S6C) and protein (Figure S6B) changes for TMBIM5 in cells treated as indicated.

(E) Representative immunoblot of WT HeLa cells transfected with scrambled siRNA (control) or siRNA targeting *AFG3L2* for 48 h. Samples were treated with the protein synthesis inhibitor cycloheximide (CHX; 10 μg/ml) and oligomycin (10 μM) and analyzed at the indicated time points (n=3 independent experiments).

**(F)** Mitochondrial membrane potential was monitored with TMRM in WT HeLa cells transiently transfected with scrambled siRNA (control) and siRNA targeting *AFG3L2* for 48 h. Fluorescent intensity was calculated relative to the value in the presence of CCCP (15 μM). Values are expressed as mean relative to control. Each measurement was performed in triplicate from 4 different preparations.
